# The ENCODE4 long-read RNA-seq collection reveals distinct classes of transcript structure diversity

**DOI:** 10.1101/2023.05.15.540865

**Authors:** Fairlie Reese, Brian Williams, Gabriela Balderrama-Gutierrez, Dana Wyman, Muhammed Hasan Çelik, Elisabeth Rebboah, Narges Rezaie, Diane Trout, Milad Razavi-Mohseni, Yunzhe Jiang, Beatrice Borsari, Samuel Morabito, Heidi Yahan Liang, Cassandra J. McGill, Sorena Rahmanian, Jasmine Sakr, Shan Jiang, Weihua Zeng, Klebea Carvalho, Annika K. Weimer, Louise A. Dionne, Ariel McShane, Karan Bedi, Shaimae I. Elhajjajy, Sean Upchurch, Jennifer Jou, Ingrid Youngworth, Idan Gabdank, Paul Sud, Otto Jolanki, J. Seth Strattan, Meenakshi S. Kagda, Michael P. Snyder, Ben C. Hitz, Jill E. Moore, Zhiping Weng, David Bennett, Laura Reinholdt, Mats Ljungman, Michael A. Beer, Mark B. Gerstein, Lior Pachter, Roderic Guigó, Barbara J. Wold, Ali Mortazavi

## Abstract

The majority of mammalian genes encode multiple transcript isoforms that result from differential promoter use, changes in exonic splicing, and alternative 3’ end choice. Detecting and quantifying transcript isoforms across tissues, cell types, and species has been extremely challenging because transcripts are much longer than the short reads normally used for RNA-seq. By contrast, long-read RNA-seq (LR-RNA-seq) gives the complete structure of most transcripts. We sequenced 264 LR-RNA-seq PacBio libraries totaling over 1 billion circular consensus reads (CCS) for 81 unique human and mouse samples. We detect at least one full-length transcript from 87.7% of annotated human protein coding genes and a total of 200,000 full-length transcripts, 40% of which have novel exon junction chains.

To capture and compute on the three sources of transcript structure diversity, we introduce a gene and transcript annotation framework that uses triplets representing the transcript start site, exon junction chain, and transcript end site of each transcript. Using triplets in a simplex representation demonstrates how promoter selection, splice pattern, and 3’ processing are deployed across human tissues, with nearly half of multi-transcript protein coding genes showing a clear bias toward one of the three diversity mechanisms. Evaluated across samples, the predominantly expressed transcript changes for 74% of protein coding genes. In evolution, the human and mouse transcriptomes are globally similar in types of transcript structure diversity, yet among individual orthologous gene pairs, more than half (57.8%) show substantial differences in mechanism of diversification in matching tissues. This initial large-scale survey of human and mouse long-read transcriptomes provides a foundation for further analyses of alternative transcript usage, and is complemented by short-read and microRNA data on the same samples and by epigenome data elsewhere in the ENCODE4 collection.

## INTRODUCTION

Most mammalian genes produce multiple distinct transcript isoforms ^1^. This transcript structure diversity is governed by promoter selection, splicing, and polyA site selection, which respectively dictate the transcript start site (TSS), exon junction chain (the unique series of exon-exon junctions used in a transcript), and transcript end site (TES, and the resulting 3’ UTR) used in the final transcript. Each of these processes is highly regulated and is subject to a different set of evolutionary pressures ^2–5^. In protein coding genes, missplicing can lead to nonfunctional transcripts by disrupting canonical reading frames or introducing premature stop codons that predispose the transcript to nonsense mediated decay (NMD). Conversely, the cellular machinery involved in promoter or polyA site selection for protein coding genes is only constrained by the need to include start and stop codons for the correct open reading frame (ORF) in the final mRNA product.

Transcript structure diversity poses challenges for both basic and preclinical biology. As computational gene prediction and manual curation efforts have identified ever more transcripts for many genes ^6,7^, a common assumption in genomics and medical genetics is that we only need to consider one or at most a handful of representative transcripts per gene such as those from the MANE (Matched Annotation from NCBI and EMBL-EBI) project ^8^. MANE transcripts are chosen with respect to their expression levels in biologically-relevant samples and sequence conservation of the coding regions, and are perfectly matched between NCBI and ENSEMBL with explicit attention to the 5’ and 3’ ends. This decision to focus on one transcript per gene was driven in part by the difficulties in transcript assembly using ESTs and short-read RNA-seq, which is the assay used for most bulk and single-cell RNA-seq experiments ^9,10^. The advent of long-read platforms heralded the promise of full-length transcript sequencing to identify expressed transcript isoforms, thus potentially bypassing the error-prone transcript assembly step ^11,12^. However, as long-read RNA-seq (LR-RNA-seq) produces more novel candidate transcripts, there is a need to find organizational principles that will allow us to cope with the diversity of transcripts observed at some gene loci in catalogs such as GENCODE ^7^, while at the same time distinguishing the genes that do not seem to undergo any alternative splicing.

Short-read RNA-seq has been the core assay for measuring gene expression in the second and third phases of the ENCODE project for all RNA biotypes, regardless of their lengths, in both human and mouse samples ^13–15^. Short-read RNA-seq has also been used by many groups to comprehensively characterize TSS usage ^16^, splicing ^17^, and TES usage ^18^, but the challenges of transcript assembly given the combinatorial nature of the problem have precluded a definitive assessment of the transcripts present. In addition to continuing Illumina-based short-read sequencing of mRNA and microRNA, the fourth phase of ENCODE (ENCODE4) adds matching LR-RNA-seq using the Pacific Biosciences Sequel 1 and 2 platforms in a set of human and mouse primary tissues and cell lines in order to identify and quantify known and novel transcript isoforms expressed across a diverse set of samples. We report the resulting ENCODE4 human and mouse transcriptome datasets. We implement a novel triplet scheme that captures essential differences in 5’ end choice, splicing, and 3’ usage, which allows us to categorize genes based on features driving their transcript structure diversity using a new software package called Cerberus. We introduce the gene structure simplex as an intuitive coordinate system for comparing transcript usage between genes and across samples. We then compare transcript usage between orthologous genes in human and mouse and identify substantial differences in transcript diversity for over half the genes.

## Results

### The ENCODE4 RNA dataset

This LR-RNA-seq study profiled 81 tissues or cell lines by using the PacBio sequencing platform on 264 human and mouse libraries that include replicate samples and multiple human tissue donors (Tables S1-2). Without consideration for the seven postnatal time-points in mouse, they represent 49 unique tissues or cell types across human and mouse (Fig. 1a, Fig. S1). In addition, we sequenced matching human short-read RNA-seq (Fig. S1c) and microRNA-seq (Fig. S2, Supplementary results) for most samples as well as for an additional 37 that were sequenced with short-read RNA-seq only. We detect the vast majority of polyA genes (those with biotype protein coding, pseudogene, or lncRNA) whether we restrict the analysis to short-read samples that have matching data in the LR-RNA-seq dataset (93.9% of GENCODE v40 polyA genes and 90.6% of protein coding genes) or if we use all of the short-read samples (Fig. 1b, Fig. S3a). 31.1% of all expressed genes are detected in most (>90%) of the samples, and 34.0% are detected more specifically (<10% of samples) (Fig. 1c, Fig. S3b).

**Figure 1.**
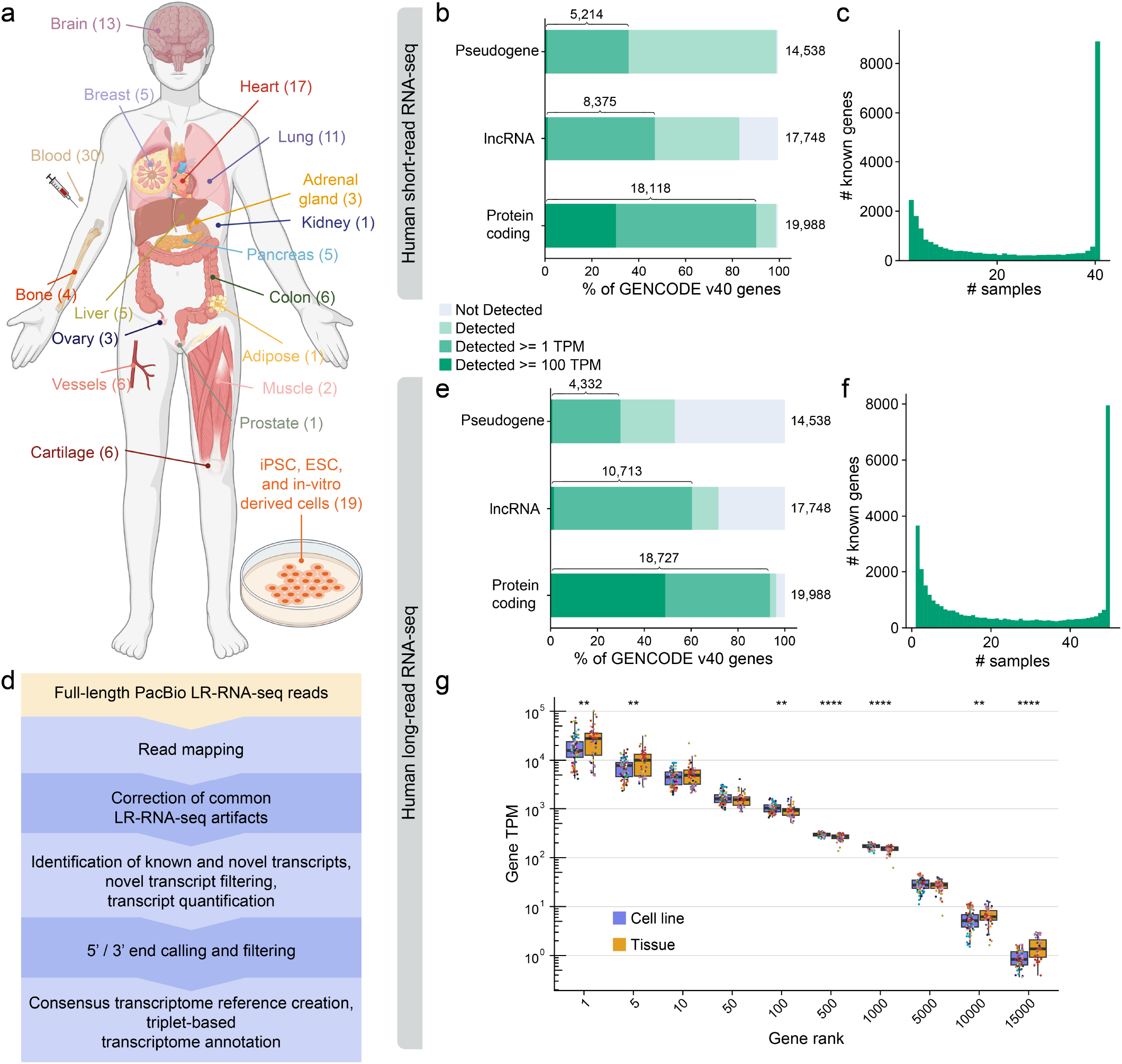
Overview of the ENCODE4 RNA datasets. a, Overview of the sampled tissues and number of libraries from each tissue in the ENCODE human LR-RNA-seq dataset. b, Percentage of GENCODE v40 polyA genes by gene biotype detected in at least one ENCODE short-read RNA-seq library from samples that match the LR-RNA-seq at > 0 TPM, >=1 TPM, and >= 100 TPM. c, Number of samples in which each GENCODE v40 gene is detected >= 1 TPM in the ENCODE short-read RNA-seq dataset from samples that match the LR-RNA-seq. d, Data processing pipeline for the LR-RNA-seq data. e, Percentage of GENCODE v40 polyA genes by gene biotype detected in at least one ENCODE human LR-RNA-seq library at > 0 TPM, >= 1 TPM, and >= 100 TPM. f, Number of samples in which each GENCODE v40 gene is detected >= 1 TPM in the ENCODE human LR-RNA-seq dataset. g, Boxplot of TPM of polyA genes at the indicated rank in each human LR-RNA-seq library. Not significant (no stars) P > 0.05; *P <= 0.05, **P <= 0.01, ***P <= 0.001, ****P <= 0.0001; Wilcoxon rank-sum test.

For each LR-RNA-seq dataset, we first mapped the reads using Minimap2^19^ and corrected non-canonical splice junctions and small indels using TranscriptClean ^20^, after which we ran TALON ^21^ and LAPA ^22^ to identify each transcript by its exon junction chain and assign each transcript a supported 5’ and 3’ end. Finally, to catalog transcript features and summarize transcript structure diversity in our datasets, we ran Cerberus, which is described below. It is important to emphasize that this pipeline (Fig. 1d) does not attempt to assemble reads, so that every reported known transcript is observed from 5’ to 3’ end in at least one read. We further required support from multiple reads for defining valid ends. Overall this is a conservative pipeline that was designed to detect and quantify robust novel and known transcripts (Materials and Methods).

Our LR-RNA-seq reads are oligo-dT primed and we therefore expect to see high detection of transcripts from polyA genes, which we define as belonging to the protein coding, lncRNA, or pseudogene GENCODE-annotated biotypes, across our datasets. Consistent with this expectation, we detect 75.9% of annotated GENCODE v40 polyA genes and 93.7% of protein coding genes at >= 1 TPM in at least one library in our human dataset (Fig. 1e). The overwhelming majority of undetected polyA genes are pseudogenes and lncRNAs, which are likely to be either lowly expressed or completely unexpressed in the tissues assayed. As expected, GO analysis of the undetected protein coding genes yielded biological processes such as smell and taste-related sensory processes that represent genes specifically expressed in tissues that we did not assay (Fig. S3c). We find that many genes are either expressed in a sample-specific manner (27.8% in <10% of samples) or are ubiquitously expressed across many samples (28.2% in >90% of samples), consistent with the short-read samples (Fig. 1f).

Transcriptionally active regions that are absent from GENCODE are candidates for novel genes. Applying conservative thresholds that included a requirement for one or more reproducible splice junctions (Supplementary methods), we found 214 novel candidate genes with at least one spliced transcript isoform expressed >= 1 TPM in human and 96 in mouse at our existing sequencing depth. Applying the same criteria to annotated polyA genes, we find 20,716 and 18,971 genes for human and mouse respectively, meaning that plausible novel genes constitute less than 1.0% in human and 0.5% in mouse. We subsequently focus analysis on transcripts from known polyA genes.

We then examined the distribution of gene expression values across our human LR-RNA-seq dataset to characterize the abundance of genes and to assess whether we would be able to measure differences in transcript abundance at our current sequencing depths. For each library, we ranked each gene by TPM and found that the most highly expressed genes have higher TPMs in primary tissue-derived libraries than cell line-derived libraries (Fig. 1g, Supplementary methods). In particular, the tissue-derived liver libraries have the most highly expressed genes at ranks 1, 5, and 10, which include ALB and FTL, as expected. We also observed that the top 1,000 genes expressed in all but one liver library are expressed >= 100 TPM and that the top 5,000 are expressed >= 10 TPM. We can therefore confidently measure major transcript expression usage with a conservative threshold of 10 TPM for at least a third of expressed genes in each sample.

From genes expressed >= 10 TPM, we are able to capture over half (54.0%) of MANE transcripts that are 9-12 kb long (Fig. S3d). Coupled with our read length profiles, we estimate that we can reliably sequence the 99.7% of annotated GENCODE v40 polyA transcripts that are <12 kb long from end-to-end if they are highly expressed (Fig. S3d-g). In mouse, we observe similar read length profiles, sample separation, and gene detection patterns (Fig. S3h-j), including detection of 84.9% of annotated GENCODE vM25 protein coding genes at >= 1 TPM (Fig. S3h). In summary, we are able to detect most of human and mouse protein coding genes in our ENCODE LR-RNA-seq datasets at similar rates to short-read RNA-seq, and our long reads are long enough to capture the vast majority of annotated polyA transcripts.

### Different sources of transcript structure diversity

We compared the transcript start sites (TSSs), exon junction chains (ECs), and transcript end sites (TESs) observed in the human LR-RNA-seq data with other prior assays of these features and with established catalogs of these features. Cerberus is designed to identify unique TSSs, ECs, and TESs from a wide variety of inputs that include LR-RNA-seq data, reference atlases, and external transcriptional assays such as CAGE, PAS-seq, and the GTEx LR-RNA-seq dataset ^18,23^ (Supplementary methods). Cerberus numbers each TSS, EC, and TES (triplet features) based on the annotation status of the transcript that it came from (e.g. the most confidently annotated will be numbered first) as well as the order in which each source was provided (Supplementary methods). Cerberus outputs genomic regions for each unique TSS and TES and a list of coordinates for each unique EC. In all cases, the gene of origin is also annotated (Fig. S4a). Using the integrated series of cataloged triplet features, Cerberus assigns a TSS, EC, and TES to each unique transcript model to create a transcript identifier of the form Gene[X,Y,Z], which we call the transcript triplet (Fig. S4b, Fig. 2a). This strategy distinguishes the structure of two different transcripts from the same gene solely on the basis of their transcript triplets. Additionally, it gives us the ability to sum up the expression of TSSs, ECs, and TESs across the transcripts they come from to enable quantification of promoter usage, EC usage, and polyA site usage respectively.

**Figure 2.**
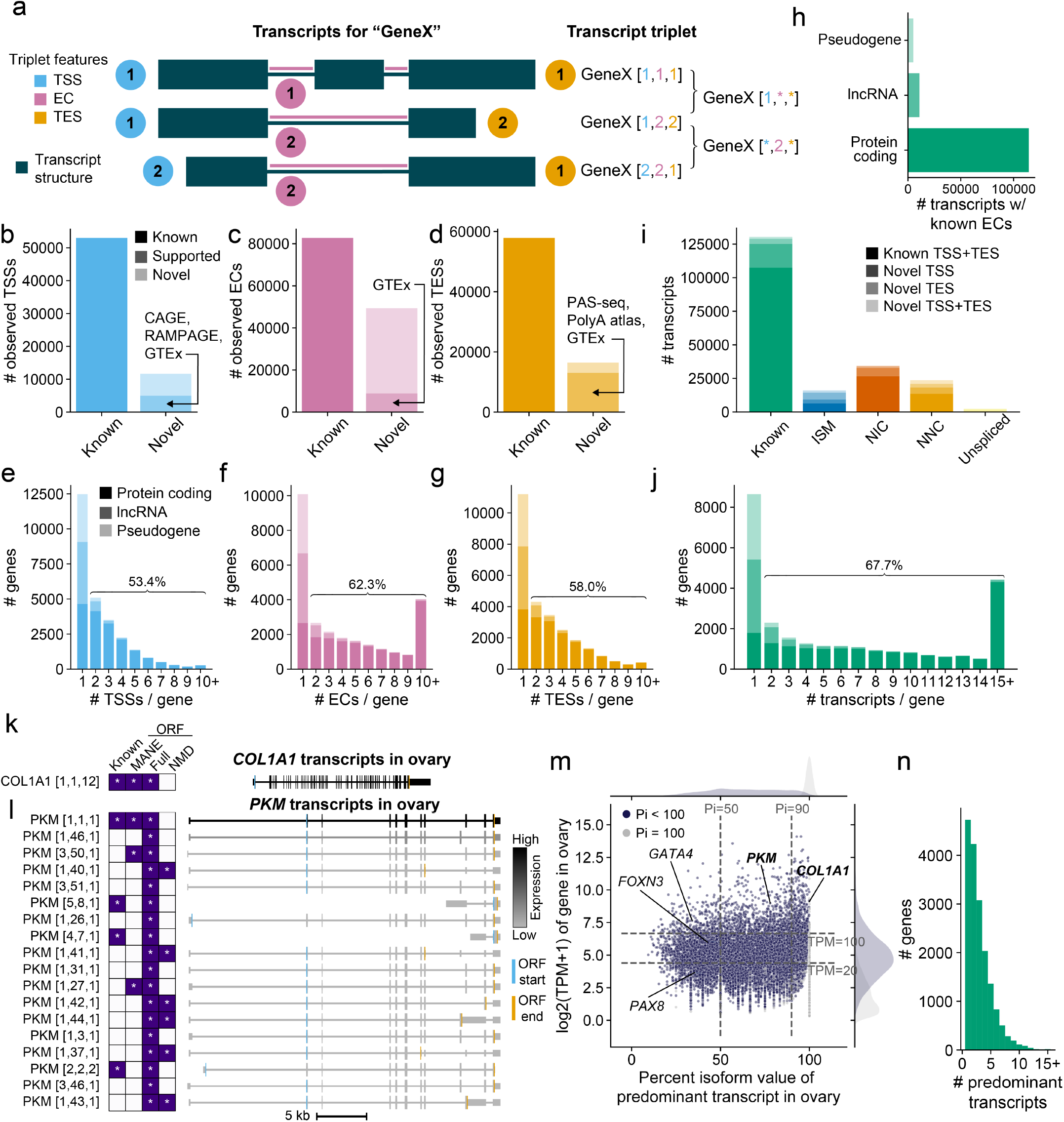
Triplet annotation of transcript structure maps diversity within and across samples. a, Representation of structure and transcript triplet naming convention for 3 different transcripts from the same gene based on the transcript start site (TSS), exon junction chain (EC), and transcript end site (TES) used. b-d, Triplet features detected >= 1 TPM in human ENCODE LR-RNA-seq from GENCODE v40 polyA genes broken out by novelty and support. Known features are annotated in GENCODE v29 or v40. Novel supported features are supported by b, CAGE or RAMPAGE c, GTEx, d, PAS-seq or the PolyA Atlas. e-g, Triplet features detected >= 1 TPM in human ENCODE LR-RNA-seq per GENCODE v40 polyA gene split by gene biotype for e, TSSs, f, ECs, g, TESs. h, Number of transcripts from GENCODE v40 polyA genes detected >= 1 TPM from human ENCODE LR-RNA-seq that have a known EC split by gene biotype. i, Novelty characterization of triplet features in each transcript detected >= 1 TPM in the human ENCODE LR-RNA-seq. j, Number of transcripts detected >= 1 TPM in human ENCODE LR-RNA-seq per GENCODE v40 polyA gene split by gene biotype. k, COL1A1 (gene expressed at 548 TPM) transcripts expressed >= 1 TPM in the ovary sample from human ENCODE LR-RNA-seq. l, PKM (gene expressed at 506 TPM) transcripts expressed >= 1 TPM in the ovary sample from human ENCODE LR-RNA-seq colored by expression level (TPM). m, Expression level of gene (TPM) versus the percent isoform (pi) value of the predominant transcript for each gene expressed >= 1 TPM from human ENCODE LR-RNA-seq in the ovary sample. Points are colored by whether or not pi = 100. n, Number of unique predominant transcripts detected >= 1 TPM across samples per gene.

We applied Cerberus to the ENCODE human LR-RNA-seq data and to annotations from GENCODE v40 and v29 to obtain transcript triplets for the transcripts present in each transcriptome. Cerberus labels each triplet feature as known if it is detected in a reference set (here defined as transcripts derived from GENCODE) of transcripts or novel if not. Additionally, using the information from the GENCODE reference transcriptomes, we assign the triplet [1,1,1] to the MANE transcript isoform for the gene, if it has one.

Altogether, we detected 206,806 transcripts expressed >= 1 TPM from polyA genes, 76,469 of which have exon junction chains unannotated in GENCODE v29 or v40. From these transcripts, we first sought to characterize the observed triplet features (expressed >= 1 TPM in at least one library from polyA genes) in our dataset (Fig. 2b-d, Fig. S5a-f). We found that 18.0% of TSSs, 37.3% of ECs, and 22.1% of TESs are novel compared to both GENCODE v29 and v40 (Fig. 2b-d). We furthermore determined whether any novel triplet features were supported by sources outside of the GENCODE reference. We used CAGE and RAMPAGE data to support TSSs, GTEx transcripts to support ECs, and PAS-seq and the PolyA Atlas regions to support the TESs (Supplementary methods). Of the novel triplet features, 42.8% of TSSs, 17.9% of ECs, and 79.0% of TESs were supported by at least one external dataset (Fig. 2b-d, Fig. S5a-c). While the intermediate transcriptome (in general transfer format; GTF) for our LR-RNA-seq dataset from LAPA has singlebase transcript ends, the majority of our Cerberus TSSs and TESs derived from the LR-RNA-seq data are 101 bp in length and 99.8% are shorter than 500 bp, which is consistent with how Cerberus extends TSSs and TESs derived from GTFs by n bp (here n=50) on either side (Fig. S4a, Fig. S5d-e, Supplementary methods).

We further annotated the novelty of ECs compared to GENCODE using the nomenclature from SQANTI ^24^. For detected ECs from polyA genes, we find that the majority (62.7%) of ECs were already annotated in either GENCODE v29 or v40. Novel ECs are primarily annotated as either NIC (16.1%; novel in catalog, defined as having a novel combination of known splice sites) or NNC (11.6%; novel not in catalog, defined as having at least one novel splice site) (Fig. S5f). Given the high external support for our triplet features, we were also able to predict CAGE and RAMPAGE support for our long-read derived TSSs both in and across cell types using logistic regression (Fig. S6, Supplementary results). In aggregate, a majority of our triplet features observed in our LR-RNA-seq data show were in prior annotations or have external support from additional assays.

We then examined the number of observed triplet features per gene. We find that most protein coding genes (89.8%) express more than one triplet feature across our dataset (Fig. 2e-g). By contrast, only 33.7% of lncRNAs and 14.4% of pseudogenes express more than 1 transcript and therefore triplet feature per gene. These biotypes exhibit far less transcript structure diversity as compared to protein coding genes. Overall, we find that our observed triplet features are individually well-supported by external annotations and assays. We also show that protein coding genes are far more likely to have more than one triplet feature than lncRNAs and pseudogenes.

### The ENCODE4 LR-RNA-seq transcriptome

Following our characterization of individual triplet features, we moved on to examining our full-length transcripts. We first note that most observed transcripts with known ECs belong to the protein coding biotype (Fig. 2h). In contrast to the genelevel analysis, transcripts with known ECs are expressed in a more sample-specific manner, with 49.0% expressed in <10% of samples and 4.4% expressed in >90% of samples (Fig. S7a, Fig. 1f). Of the remaining protein coding transcripts with novel ECs, 53.0% are predicted to have complete ORFs which are not subject to nonsense mediated decay (Supplementary methods). Examining detected transcripts in our dataset based on the novelty of each constituent triplet feature, we find that 52.0% of observed transcripts have each of their triplet features annotated, and that more transcripts contain novel TSSs than novel TESs (Fig. 2i). Consistent with our observation that protein coding genes generally have more than one triplet feature per gene compared to the other polyA biotypes, we find that most protein coding genes also have more than one transcript per gene (Fig. 2j, e-g). We investigated the extent that lower expression levels of lncRNAs contribute to their overall lower transcript diversity compared to protein coding genes. We found that the 137 lncRNAs expressed >100 TPM in one or more samples have the same median number of expressed transcripts per gene as the 8,436 protein coding genes at the same expression level (median = 7) (Fig. S7b, Supplementary methods). Therefore the lower reported overall diversity of lncRNAs is due to a combination of their lower expression levels and our sequencing depth.

We compared the number of TSSs and TESs that are detected per EC across the observed transcripts from GENCODE v40 versus in our observed transcripts. We found that in GENCODE v40, each multiexonic EC has a maximum of 3 TSSs or TESs across the polyA transcripts and that the overwhelming majority of ECs are only annotated with 1 TSS and TES (99.7% and 99.4% respectively) (Fig. S7c-d). In contrast, our strategy of transcriptome annotation yields a substantial increase in the number of distinct TSSs and TESs observed per EC, which more accurately reflects the biology of the coordination of promoter choice, polyA site selection, and splicing across the diverse samples in our dataset (Fig. S7e-f). The effect of this increase in annotated TSSs and TESs is also apparent when analyzing our transcripts using traditional alternative splicing event detection methods, which are not written to consider the more subtle differences in transcript structure at the 5’ and 3’ end (Fig. S8, Supplementary results).

### Predominant transcript structure differs across tissues and cell types

Different multiexonic genes with similar expression levels within the same sample can exhibit vastly different levels of transcript structure diversity. For instance, the genes COL1A1 and PKM have a high number of exons (60 and 47 exons, respectively across our entire human dataset) and are highly expressed in ovary (548 and 506 TPM respectively). Yet, we detect only one 6.9 kb long transcript for COL1A1 (Fig. 2k) whereas we detect 18 transcript isoforms that vary on the basis of their TSSs, ECs, and TESs for PKM (Fig. 2l).

We then asked what fraction of overall gene expression is accounted for by the predominant transcript, which is the most highly expressed transcript for a gene in a given sample. Comparing the TPM of genes expressed in ovary to the the percentage of reads from a gene that come from that transcript (pi -percent isoform) ^25^ of the predominant transcript, we find that 19.5% of protein coding genes expressed >100 TPM have a predominant transcript that accounts for less than 50% of the reads, and therefore are highly expressed with high transcript structure diversity. Conversely, 26.8% of protein coding genes are expressed >100 TPM and have a predominant transcript that accounts for more than 90% of the expression of the gene (Fig. 2m). Globally, we generated a catalog of predominant transcripts for each sample. The median number of predominant transcripts per protein coding gene across samples was 2, and that 73.0% of protein coding genes have more than one predominant transcript across the samples surveyed (Fig. 2n). Thus, the majority of human protein coding genes use a different predominant transcript in at least one condition represented in our sample collection.

### Quantifying transcript structure diversity across samples using gene triplets and the gene structure simplex

We developed a framework to systematically characterize and quantify the diversity between the detected transcripts from each gene by computing a summary gene triplet, which is related to but distinct from transcript triplets. For each set of transcripts from a given gene, we count the number of unique TSSs, ECs, and TESs (Fig. 3a, Fig. S9). As the number of exon junction chains is naturally linked to the number of alternative TSSs or TESs (for instance, a new TSS with a different splice donor will lead to a novel EC regardless of similarities in downstream splicing), we calculate the splicing ratio as 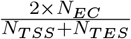 to more fairly assess the contribution of ECs to transcript diversity in each gene (Fig. 3a, Fig. S9). We then compute the proportion of transcript diversity that arises from each source of variation: alternative TSS usage, alternative TES usage, or internal splicing (Fig. 3a). Representing these numbers as proportions allows us to plot them as coordinates in a two-dimensional gene structure simplex (Fig. 3b, Fig. S9). This enables us to visualize how transcripts from a gene typically differ from one another and categorize genes based on their primary driver of transcript structure diversity. Genes with a high proportion of transcripts characterized by alternative TSS usage (>0.5) will fall into the TSS-high sector of the simplex, those with a high proportion of transcripts characterized by alternative TES usage (>0.5) will fall into the TES-high sector of the simplex, and those with a high proportion of transcripts characterized by internal splicing (>0.5) will fall into the splicing-high portion of the simplex. Genes with more than one transcript that do not display a strong preference for one mode over the other lie in the mixed sector, and genes with just one transcript are in the center of the simplex, henceforth the simple sector (Fig. 3a-b, Fig. S9, Supplementary methods).

**Figure 3.**
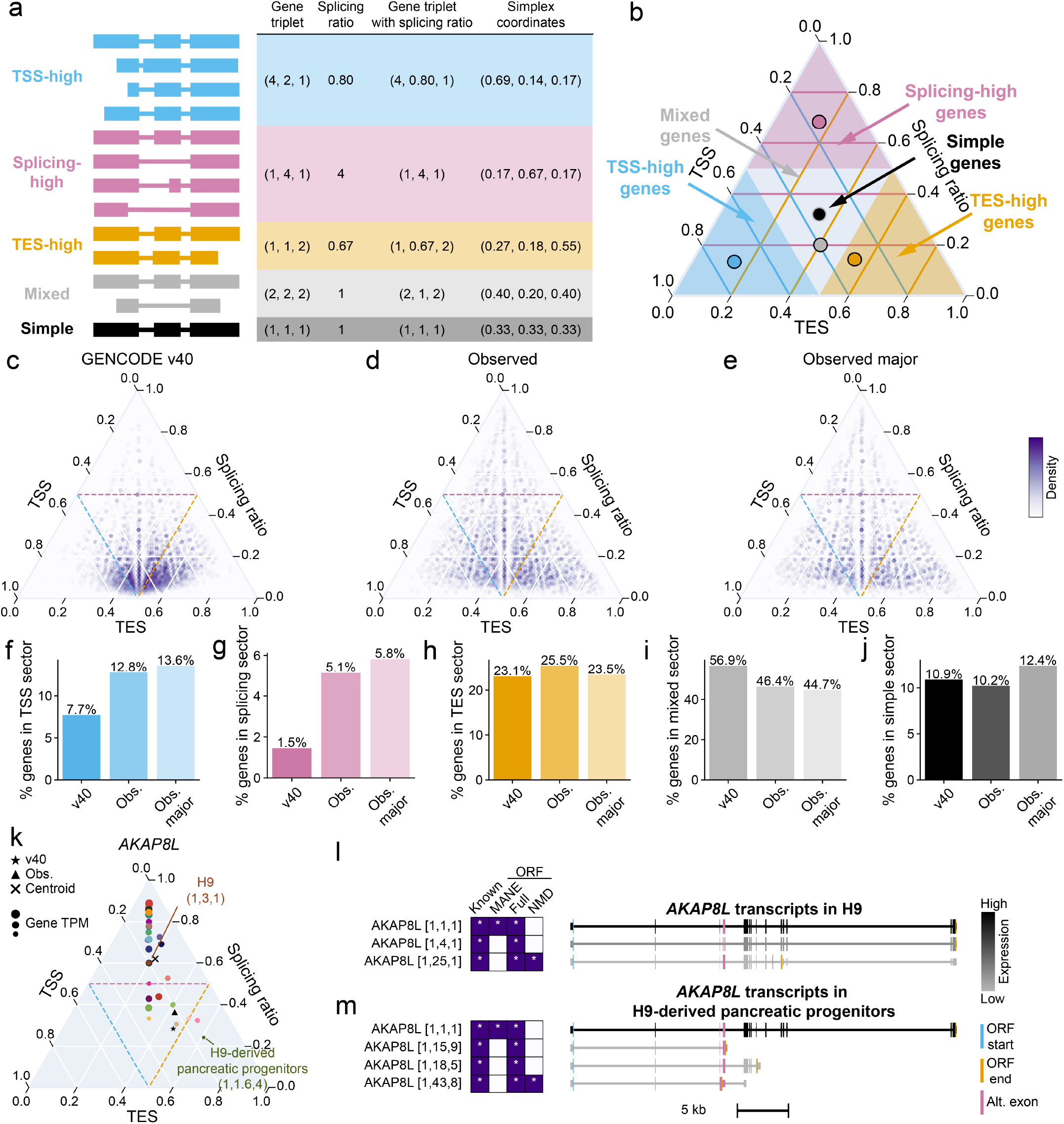
The gene structure simplex represents distinct modes of transcript structure diversity across genes and samples. a, Transcripts for 5 model genes; 1 of each sector (TSS-high, splicing-high, TES-high, mixed, and simple). Table shows the gene triplet, splicing ratio gene triplet, and simplex coordinates that correspond to each toy gene. b, Layout of the gene structure simplex with the genes from a, plotted based on their simplex coordinates. Proportion of TSS usage is the blue axis (left), proportion of TES is the orange axis (bottom), and proportion of splicing ratio is the pink axis (right). Regions of the simplex are colored and labeled based on their sector category (TSS-high, splicing-high, TES-high). Gene triplets that land in each sector are assigned the concordant sector category. c-e, Gene structure simplices for the transcripts from protein coding genes that are c, annotated in GENCODE v40 where the parent gene is also detected in our human LR-RNA-seq dataset, d, the observed set of transcripts, those detected >= 1 TPM in the human ENCODE LR-RNA-seq dataset, e, the observed major set of transcripts, the union of major transcripts from each sample detected >= 1 TPM in the human ENCODE LR-RNA-seq dataset. f-j, Proportion of genes from the GENCODE v40, observed, and observed major sets that fall into the f, TSS-high sector, g, splicing-high sector, h, TES-high sector, i, mixed sector, j, simple sector. k, Gene structure simplex for AKAP8L. Gene triplets with splicing ratio for H9 and H9-derived pancreatic progenitors labeled. Simplex coordinates for the GENCODE v40, observed set, and centroid of the samples also shown for AKAP8L. l-m, Transcripts of AKAP8L expressed >= 1 TPM in l, H9 m, H9-derived pancreatic progenitors colored by expression level in TPM. Alternative exons that differ between transcripts are colored pink.

We first used the gene structure simplex to compare different transcriptomes. We computed gene triplets for protein coding genes for the following transcriptomes: GENCODE v40 transcripts from genes we detect in our LR-RNA-seq dataset; observed transcripts in our LR-RNA-seq dataset (observed); and the union of detected major transcripts (observed major), which we define as the set of most highly expressed transcripts per gene in a sample that are cumulatively responsible for over 90% of that gene’s expression in any of our LR-RNA-seq samples (Fig. 3c-j, Supplementary methods). The observed and observed major gene triplets describe the diversity of transcription in each gene across all samples in the dataset. Unsurprisingly, GENCODE genes show less density in the TSS or TES sectors of the simplex, largely because the main focus of GENCODE is to annotate unique ECs rather than 5’ or 3’ ends. This causes a concomitant drop in diversity in these sectors in GENCODE compared to the observed and observed major transcripts (Fig. 3f, h). Interestingly, there is also a distinct enrichment of genes that occupy the splicing-high portion of the simplex in our observed set compared to GENCODE (Fig. 3g). When considering the observed major transcripts, we see an increase in the percentage of genes in the TSS and splicing-high over the set of all transcripts detected in our entire LR-RNA-seq dataset, but a decrease in the TES-high sector (Fig. 3f-h). Overall, we compared gene triplets for transcripts as annotated by GENCODE and observed in our LR-RNA-seq dataset and found higher proportions of genes with high TSS and splicing diversity as compared to GENCODE.

### Calculating sample-level gene triplets identifies genes that show distinct transcript structure diversity across samples

The observed gene triplets represent the aggregate repertoire of triplet features for each gene globally across our entire LR-RNA-seq dataset. However, the overall transcript structure diversity of a gene does not necessarily reflect the transcript structure diversity of a gene within a given sample. Therefore, we computed gene triplets for each sample in our dataset using all detected transcripts in each sample (sample-level gene triplets) or just the major transcripts in each sample (sample-level major gene triplets). These gene triplets can also be visualized on the gene structure simplex where each point represents the gene triplet associated with a different sample (Fig. 3k).

In order to find genes that display heterogeneous transcript structure diversity across unique biological contexts, we computed the average coordinate (centroid) for each gene from the sample-level gene triplets and calculated the distance between it and each sample-level gene triplet (Supplementary methods). 2,892 unique genes had a distance z-score >3 in at least one sample and therefore demonstrate dissimilar transcript structure diversity from the average. One such example is AKAP8L in the H9-derived pancreatic progenitors (z-score: 5.23). AKAP8L can bind both DNA and RNA in the nucleus and has been shown to have functional differences on the protein level resulting from alternate transcript choice ^26,27^. In our data, transcripts of this gene generally differ in terms of the EC or TES choice, but this behavior differs from sample to sample (Fig. 3k). For example, transcripts of AKAP8L differ only in their ECs in H9 embryonic stem cells, whereas transcripts differ in their ECs and TESs in the H9-derived pancreatic progenitors (Fig. 3l-m).

We also compared our sample-level gene triplets to the observed gene triplets to understand how transcript structure diversity differs globally versus within samples (Fig. 3d, Fig. 2j, Fig. S10a-h). First, we simply counted the number of triplet features or transcripts per gene and found that while most genes have more than one triplet feature or transcript globally (Fig. 2e-g, j), on the sample level, most genes have far fewer triplet features and transcripts; with a particularly pronounced difference for the TSS (Fig. S10a-d). We found that the distributions of triplet features overall and in each sample were significantly different from one another (twosided KS test) (Fig. S10e-l, Supplementary methods).

To determine how transcript structure diversity for each gene changes from the global to sample level using the gene structure framework, we computed distances between the global observed gene triplets for non-simple genes to sample-level gene triplet centroids (Supplementary methods). We find that 3.2% of tested genes have a distance z-score >2 between their observed and sample-level centroid gene triplets. In support of our analysis on the individual triplet feature level, we find that 94.8% of genes from the TSS-high sector in the observed set do not share this sector with their sample-level centroid, indicating that genes with a large number of promoters typically use them in a sample-specific manner. ACTA1, a gene that encodes for an actin protein ^28^, is the gene with the highest distance between observed and sample-level centroid. Its observed gene triplet is (1,18,1) and therefore splicing-high. However, in most samples where ACTA1 is expressed, it has only one transcript isoform (Fig. S10m). This drives the sample-level centroid behavior into the mixed sector (Fig. S10n). In contrast, in heart and muscle ACTA1 expresses 18 and 15 transcripts respectively, which all differ on the basis of their ECs (Fig. S10m-o). This illustrates how the gene structure framework can be used to highlight differences between sample-specific and global transcript structure diversity, and also shows that individual genes are substantially different.

### Sample-specific and global changes in predominant and major transcript isoform usage

Nevertheless, the transcript structure diversity pattern for the majority of genes is consistent across samples where they are expressed at substantial levels. Elastin (ELN), which is an important component of the extracellular matrix ^29^, is the gene with the greatest number of detected transcripts in our dataset (283 in total). We find that in most samples, distinct transcripts of ELN are characterized by different ECs (Fig. 4a). For example, in lung, ELN has 32 major transcripts with 21 different ECs, but in 31 of its major transcripts, uses only one TSS and two TESs (Fig. 4b). By contrast, the four transcripts from the transcription factor CTCF expressed in lung use three TSSs but only one TES (Fig. 4c-d).

**Figure 4.**
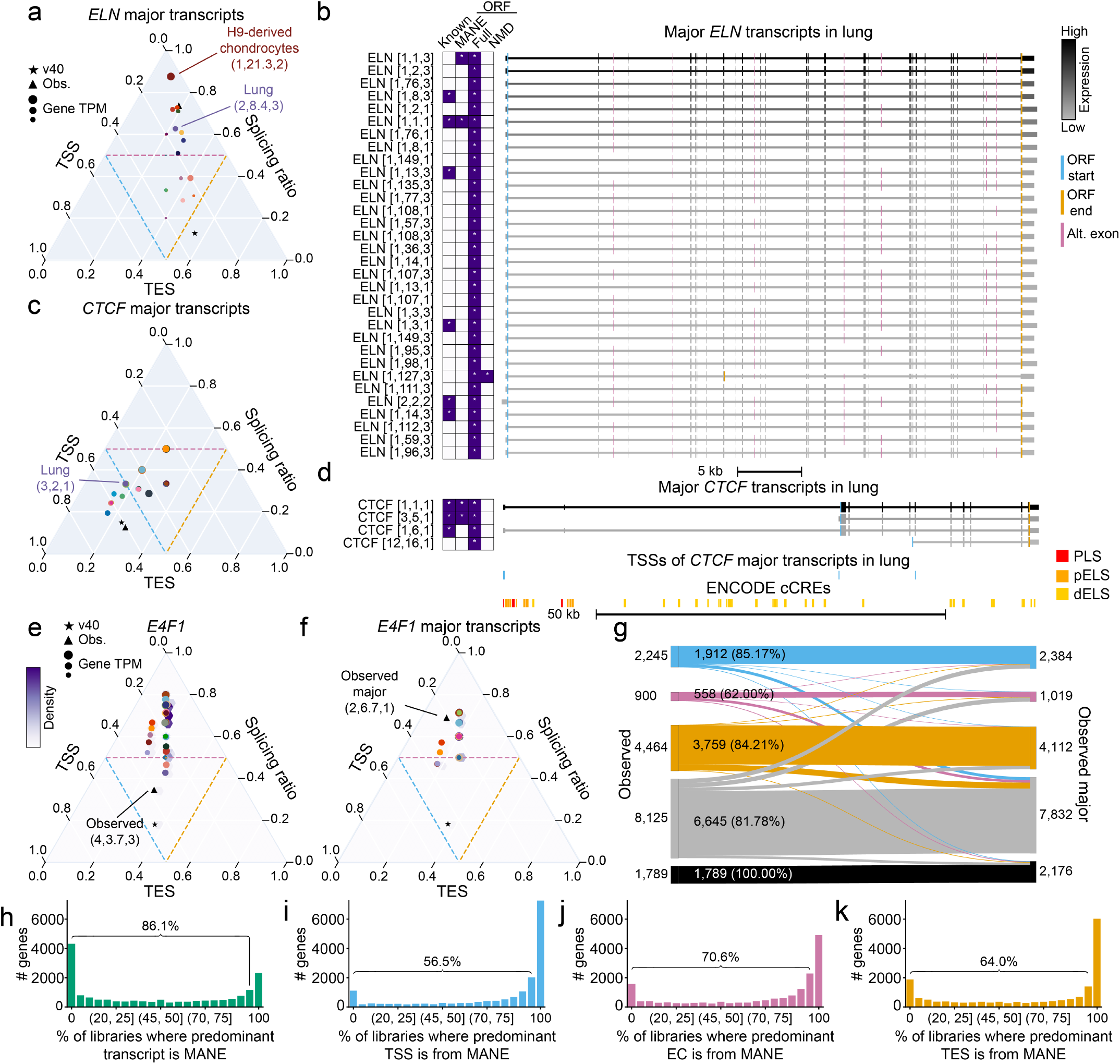
Sample-specific and global changes in predominant and major transcript isoform usage. a, Gene structure simplex for major transcripts of ELN. Gene triplets with splicing ratio for lung and H9-derived chondrocytes labeled. Simplex coordinates for the GENCODE v40 and observed major set are labeled. b, Major transcripts of ELN expressed >= 1 TPM in lung colored by expression level in TPM. Alternative exons that differ between transcripts are colored pink. c, Gene structure simplex for major transcripts of CTCF. Gene triplets with splicing ratio for lung labeled. Simplex coordinates for the GENCODE v40 and observed major set are labeled. d, From top to bottom: Major transcripts of CTCF expressed >= 1 TPM in lung, TSSs of CTCF major transcripts expressed >= 1 TPM in lung, ENCODE cCREs colored by type. e, Gene structure simplex for E4F1. Gene triplet with splicing ratio for observed E4F1 transcripts labeled. Simplex coordinates for the GENCODE v40 and observed set also shown for E4F1. f, Gene structure simplex for major transcripts of E4F1. Gene triplet with splicing ratio for observed major E4F1 transcripts labeled. Simplex coordinates for the GENCODE v40 and observed major set also shown for E4F1. g, Sector assignment change and conservation for protein coding genes in the human ENCODE LR-RNA-seq dataset between the observed set of gene triplets (left) and the observed major set of gene triplets (right). Percent of genes with the same sector between both sets labeled in the middle. h-k, Percentage of libraries where a gene with an annotated MANE transcript is expressed and the MANE h, transcript i, TSS j, EC k, TES is the predominant transcript or triplet feature.

While the observed gene triplets for a gene represent the overall transcript structure diversity, the observed major gene triplets capture diversity of the most highly expressed transcripts in each sample. We computed the distances between the observed and observed major simplex coordinates for protein coding genes. The transcription factor E4F1 ^30^ has a high distance between the observed and observed major gene triplets, which corresponds to a change from the mixed to splicing-high sector (Fig. 4e-f). This sector change is driven by the use of fewer TSSs and TESs in major transcripts. Overall, 83.7% of protein coding genes retain their sectors between our observed and observed major triplets, while 4.8% genes in the mixed sector move to one of the three corners of the simplex (TSS, splicing, or TES-high) (Fig. 4g). Thus, the differences between the observed and observed major gene triplets in a subset of genes can be substantial.

One criterion for the identification of MANE transcripts is how highly expressed the transcript is compared to others ^8^. Therefore, we assessed how frequently the MANE transcript was the predominant one in each of our LR-RNA-seq libraries. Limiting ourselves to only the genes that have annotated MANE transcripts in GENCODE v40, we found that 64.1% of genes have a non-MANE predominant transcript in at least 80% of the libraries where the gene is expressed (Fig. 4h). At the individual triplet feature level, 30.8% of TSSs, 40.9% of ECs, and 45.2% of TESs have a non-MANE predominant feature in at least 80% of libraries (Fig. 4i-k). Therefore, though the MANE transcript typically is the most highly expressed transcript in a library, most genes with MANE transcripts have some libraries where this is not the case. For non-MANE predominant transcripts, only 17.0% were predicted to have the same ORF as the MANE transcript. Furthermore, 62.1% of non-MANE predominant transcripts are predicted to encode for a full ORF that does not undergo NMD. These results indicate that in many cases, the alternative predominant transcript in a sample likely encodes for a distinct, functional protein. The genes where the MANE transcript and triplet features are frequently not the predominant one represent loci that would suffer more from restricting analyses to only a single transcript isoform.

For a subset of gene / library combinations where the MANE transcript or feature was not the predominant one, the MANE transcript or feature was still expressed, albeit at a lower level. For these gene / library combinations, we compared the expression of the predominant transcript to the MANE one (Fig. S11a-d). We found that for predominant transcripts or triplet features expressed <30 TPM, the MANE counterpart was expressed at a comparable level. By contrast, for the opposite situation, where the MANE transcript or triplet feature was the predominant one, we found that the secondary transcript was not expressed at a similar level (Fig. S11e-h). Overall, for most gene / library combinations, the MANE transcript or triplet feature is the predominant one (Fig. S11i-l).

### Comparing transcript structure diversity between species

We ran Cerberus on the ENCODE4 mouse LR-RNA-seq dataset to calculate transcript and gene triplets to enable comparison of transcript structure diversity between the two species. Compared to human GENCODE v40, GENCODE vM25 genes are less enriched in the TSS, splicing, and TES-high sectors (Fig. 3f-j, Fig. 5a-e, Fig. S12a-c). For the mouse observed and observed major gene triplets, we see relatively similar percentages of genes in each sector (Fig. 5a-e, Fig. S12a-c). We found fewer predominant transcripts across samples per protein coding gene in mouse than in human, which is expected due to the overall lower number of tissues in our mouse data, with a median of 2 predominant transcripts per gene and 57.7% of protein coding genes with more than one (Fig. S12d). Furthermore, we observe that 54.5% of protein coding transcripts with novel ECs are predicted to encode for full ORFs without NMD. Thus, the two transcriptomes have similar distributions of genes in our gene structure simplex.

**Figure 5.**
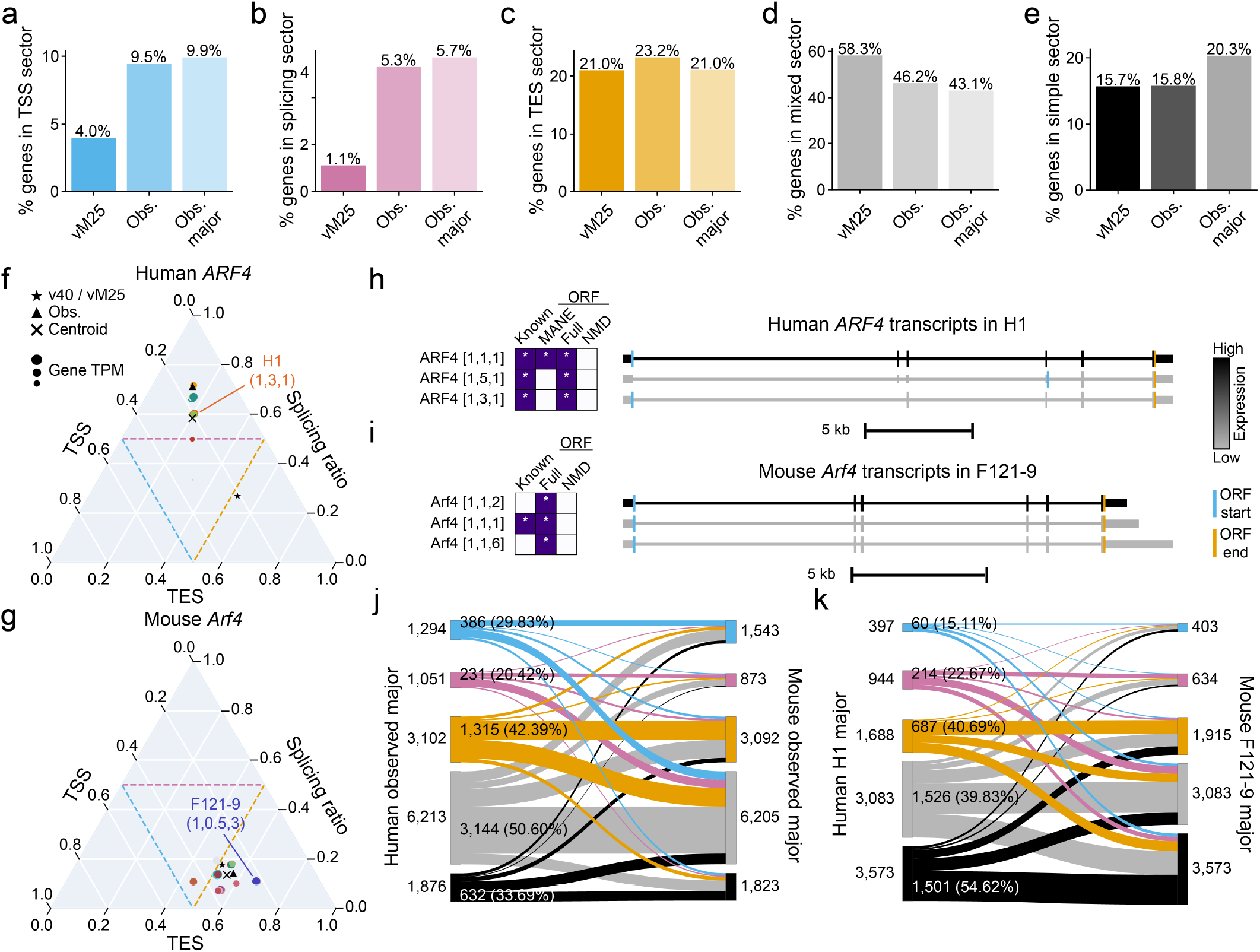
Conservation of gene triplets from human and mouse. a-e, Proportion of genes from the GENCODE vM25, observed, and observed major sets that fall into the a, TSS-high sector, b, splicing-high sector, c, TES-high sector, d, mixed sector, e, simple sector. f, Gene structure simplex for ARF4 in human. Gene triplet with splicing ratio for ARF4 transcripts in H1 labeled. Simplex coordinates for the GENCODE v40, sample-level centroid, and observed set also shown for ARF4. g, Gene structure simplex for Arf4 in mouse. Gene triplet with splicing ratio for Arf4 transcripts in F121-9 labeled. Simplex coordinates for the GENCODE v40, sample-level centroid, and observed set also shown for Arf4. h, Transcripts of ARF4 expressed >= 1 TPM in human H1 sample colored by expression level in TPM. i, Transcripts of Arf4 expressed >= 1 TPM in mouse F121-9 sample colored by expression level in TPM. j, Sector assignment change and conservation for orthologous protein coding genes between the observed major human set of gene triplets (left) and the observed major mouse set of gene triplets (right). Percent of genes with the same sector between both sets labeled in the middle. k, Sector assignment change and conservation for orthologous protein coding genes between the sample-level H1 major human set of gene triplets (left) and the sample-level F121-9 major mouse set of gene triplets (right). Percent of genes with the same sector between both sets labeled in the middle.

In order to make gene-level comparison for orthologous genes in both species, we subset the human samples on those that are the most similar to the mouse samples and computed “mouse matched” observed, observed major, and sample-level gene triplets (Supplementary methods, Table S1-2). We computed the sample-level centroids for each gene in both species and computed the distance between each pair of 1:1 orthologs. Of the 13,536 orthologous genes, 4.3% have a distance z-score >2 between the species and therefore exhibit substantial changes in transcript structure diversity between the species. One of these is ADP-Ribosylation Factor 4 (ARF4), which is the most divergent member of the ARF4 family ^31^. Human ARF4 sample-level gene triplets are nearly always splicing-high whereas mouse Arf4 sample-level gene triplets are mainly TES-high (Fig. 5f-g). We examined the ARF4 / Arf4 transcripts expressed in matching embryonic stem cell samples (H1 in human and F121-9 in mouse) and found that, despite the homologous samples, all 3 of the expressed human transcripts use the same TSS and TES but differ in the ECs whereas all 3 expressed mouse transcripts use the same TSS and EC but differ at the TES (Fig. 5h-i). We find globally that when comparing the observed major gene triplets between human and mouse, only 42.2% of genes have the same sector in human and mouse (Fig. 5j). This result holds even when restricting ourselves to comparing human tissues with adult mouse samples or just a comparison between human and mouse embryonic stem cells (Fig. 5k). Thus, we find substantial differences in splicing diversity for orthologous genes between human and mouse.

## Discussion

The ENCODE4 LR-RNA-seq dataset is the first large-scale, cross-species survey of transcript structure diversity using full-length cDNA sequencing on long-read platforms. We identify and quantify known and novel transcripts in a broad and diverse set of samples with uniformly processed data and annotations available at the ENCODE portal. A new framework was introduced for categorizing transcript structure diversity based on their exon junction chains and ends using gene and transcript triplets, which allowed us to use the gene structure simplex to visualize and compare gene triplets between samples and across species. The results showed a full range of transcript structure diversity across the transcriptome, based on promoter, internal splicing, and polyA site choice. As expected, the existing gene annotation catalogs such as GENCODE have successfully captured individual features such as TSSs and exons. However, GENCODE annotated full-length transcripts only represent a subset of the TSS, EC, and TES combinations that we observe using our conservative pipeline that requires full end-to-end support in a single read and support from multiple reads for defining ends. From the human LR-RNA-seq quantification, we found more than one predominant transcript across samples for 73.0% of genes, which is in contrast with prior reports ^32,33^. We also found that for a substantial number of genes, transcript structure diversity and major transcript usage for the same gene differs between tissues, samples, and developmental timepoints. The majority of genes had at least one library where the MANE transcript is not the predominant transcript. This could confound analyses such as variant effect prediction in which it is common practice to consider only one transcript per gene. Finally, we found that transcript structure diversity behavior differs quite strikingly between human and mouse on a gene-by-gene basis. In matching samples, the dominant source of transcript structure diversity differed for more than half of orthologous protein coding genes.

Our data and framework provide a foundation for further analyses such as the functional impacts of alternative 5’ and 3’ ends, RNA modifications, RNA binding protein function, allele-specific expression, and transcript half-life. Together with the accompanying tissue and cell type annotations, this constitutes a transcript-level reference atlas that is structured appropriately for integration of future single-cell long-read analysis. Exploration using the gene structure simplex analysis will yield additional genes showing sample specific variance compared to their average behavior when extended to new tissues, differentiation time courses, or disease samples. The triplet annotation scheme for transcripts, based on mechanistically distinct transcript features, organizes and simplifies high-level analysis of transcripts from the same gene. We find it to be a useful and commonsense improvement over arbitrary transcript IDs and we expect it to be widely applicable to transcriptomes of any organism that uses regulated alternative splicing, promoter choice, or 3’ end selection.

Our annotations are consistent and extensive, yet they have several limitations. With the current sample preparation protocol and depth of sequencing, we reliably detect transcripts that are expressed above a minimum expression level of 1 TPM and are less than 10 kb long. While 99.3% of GENCODE v40 polyA transcripts are less than 10 kb long, we are undoubtedly underrepresenting transcripts that are at the long end of the distribution, especially when they are expressed at low levels or in rare cell types within a tissue. RNA integrity differences between the human cell lines and mouse tissues, both of which produce very high quality RNA compared to human postmortem tissues, are expected to affect our results, because we have focused on full-length transcript sequencing rather than read assembly. Our imposition of minimum expression and inherent length limitations could also lead to lower sensitivity of splicing diversity in lncRNAs, which have thus far generated staggering transcript structure complexity when sequenced after enrichment capture ^34^. We also had lower detection of pseudogenes, and we hypothesize that the PacBio platform’s accuracy and read length reduce the multimapping errors typical of short reads, especially for pseudogenes of highly expressed genes.

Within these boundaries outlined above, we were able to assess the sources and specifics of transcript structure diversity for major transcripts of most protein coding genes. Nearly all studies that have examined alternative splicing have emphasized transcript isoform multiplicity per gene. Also as expected, studies that have applied more permissive processing pipelines, used transcript assembly, or focused on nuclear RNA typically find evidence for far more RNA transcripts, especially at lower expression ranges ^24,35–38^. Assigning biological functions to new transcripts from our collection or any other contemporary study is a major challenge for the field. Unlike DNA replication’s elaborate mechanisms to ensure fidelity, the three major processes of RNA biogenesis mapped here are understood to operate with less stringent fidelity, and though it has long been debated, we consider evidence for the existence of a new transcript isoform simply makes it a candidate of interest for a protein coding, precursor, or regulatory function.

The range of regulation used by different genes was illuminating. COL1A1, a complex gene in terms of number of exons, exhibited minimal transcript structure diversity in spite of high expression versus other genes of similar expression levels, such as PKM with its many transcripts resulting from all three mechanisms. This implies that transcript structure diversity is a property of the gene that has been optimized in evolution. This has major implications for evaluating the functions of regulatory factors such as PAX6, which has 81 transcripts in GENCODE v40, and 33 transcripts in our dataset. Conventional gene-level short-read RNA-seq profiling is likely obscuring important distinctions in transcript usage. While not every one of these transcripts leads to a difference in the protein product, changes at the 5’ and 3’ end are likely to alter the regulation of those transcripts. The incorporation of transcript usage as well as its regulation within the framework of gene regulatory networks, where appropriate, is a major challenge going forward.

Considering transcript structure diversity as a fundamental, tunable property of gene function, the mouse-human comparative results were the most surprising to us. In genomics and in the wider biology community we often use orthology of mouse and human genes to predict and interpret gene function in vivo, including many uses of mice as mammalian models for both basic and preclinical purposes. The differences in transcript structure diversity that surfaced when we compared matching tissues from human and mouse suggests that this diversity is rapidly evolving on a per gene basis, even between primates and rodents. This is, however, consistent with prior observations of a large population of rapidly evolving candidate cis-regulatory elements ^39^. The results presented here provide a roadmap for evaluating the evolution of transcript structure diversity across species and impetus to focus on it, especially for genes with substantial differences that would affect interpretation of existing animal models and expectations for humanized gene-locus mouse models. It is hard to underestimate the need for better methods to test the functional significance of different transcript isoforms.

## ACKNOWLEDGEMENTS

We thank the UCI GRTH for sequencing PacBio LR-RNA-seq libraries and the Caltech Jacobs Genetics and Genomics Laboratory for sequencing the Illumina mRNA RNA-seq libraries. We thank Gloria Sheynkman for guidance on protein prediction analyses.

A.M., B.J.W., L.R., D.B. were supported by UM1HG009443. B.J.W. was also supported by the Caltech Beckman Institute BIFGRC. D.B. was also supported by P30AG10161, P30AG72975, R01AG15819, R01AG17917, U01AG46152, U01AG61356. Z.W., M.B.G., R.G., and the members of the ENCODE DAC were supported by U24HG009446. M.L. was supported by UM1HG009382. M.B. was supported by R01HG012367 and U01HG009380. B.H. and the members of the ENCODE DCC were supported by U24HG009397.

## AUTHOR CONTRIBUTIONS

B.W., G.B., E.R., H.Y.L., C.J.M., S.R., S.J., W.Z., K.C., A.K.W., and L.A.D. did the experimental work. F.R., D.W., M.H.C., E.R., N.R., D.T., S.R., J.S., S.U., J.J., I.Y., I.G., P.S., O.J., J.S.S., M.S.K., and B.C.H. developed the data processing pipelines. F.R., E.R., N.R., D.T., M.R., Y.J., B.B., S.M., A. McShane, K.B., and S.I.E. performed data analysis. F.R., B.W., E.R., N.R., M.R., Y.J., B.B., and A. Mortazavi wrote the manuscript. All other authors provided project input and edited the manuscript.

## DATA AVAILABILITY

- Human LR-RNA-seq processed data / processing pipeline
- Human LR-RNA-seq datasets
- [Mouse LR-RNA-seq processed data / processing pipeline](https://www.encodeproject.org/annotations/ENCSR110KDI/)
- Mouse LR-RNA-seq datasets
- Human short-read RNA-seq datasets
- Human microRNA-seq datasets

## CODE AVAILABILITY

- Data processing and figure generation code
- Cerberus

## SUPPLEMENTARY RESULTS

### The ENCODE4 microRNA-seq dataset

We sequenced 254 human samples with microRNA-seq, which specifically captures mature (21-25 bp) microRNAs. We mapped our microRNA-seq data to pre-microRNA sequences in GENCODE v29. We detected 1,130 microRNAs at CPM >= 2 across the full dataset (Fig. S2a). Overall, microRNAs are more sample-specific than known genes in both short and long-read RNA-seq (Fig. S2a, Fig. 1c, f, Fig. S3b). PCA of all microRNA samples shows separation of brain samples from other tissues and cell lines by PC1, while cell lines are separated by both PC1 and PC2 (Fig. S2b). Comparison of microRNA detection between tissue types and cell lines reveals that brain samples have the least diversity when compared to the full set of GENCODE v29 microRNAs (Fig. S2c), yet the most microRNAs expressed at >=2 CPM (Fig. S2d). This indicates that a core set of microRNAs are expressed in the brain. Their high tissue-specific expression may be driving the clustering of brain samples apart from non-brain tissues and cell lines, which overlap slightly (Fig. S2b). Comparison of the overlap of detected microRNAs across the sample biotypes reveals that more microRNAs overlap between brain and cell lines than between brain and non-brain tissues (Fig. S2e). Of the 80 shared microRNAs between brain and cell lines, most (53) are expressed in neuronal and glial derived cells.

### Machine learning models predict the support for long-read TSS peaks by other TSS-annotating assays and in a cross-cell type manner

We sought to identify a set of high-confidence TSS regions from our observed LR-RNA-seq TSSs using multiple orthogonal TSS assays such as RAMPAGE and CAGE ^40,41^. However, matching data from external assays is only available for a few samples, such as our ENCODE tier 1 cell lines GM12878 and K562. Therefore, we wanted to predict the external support for our observed LR-RNA-seq TSSs. The majority of our observed TSS regions are supported by these external assays (Fig. S6a-b, Fig. 2b, Fig. S5a). We used a simple logistic regression model that incorporates expression, DNase-Hypersensitivity (DHS) ^42^, and length of our LR-RNA-seq observed TSSs (Supplementary methods, Fig. S6c-d). Models trained and tested on one experiment each from GM12878 and K562 TSS regions were able to predict whether an LR-RNA-seq TSS region was also supported by RAMPAGE or CAGE assays, with AUROC values as high as 0.95 for Cerberus and 0.98 for LAPA-annotated peaks in the same cell type (Fig. S6e), which is expected given that LAPA regions are narrower than Cerberus regions. Models trained on one cell type can also be used to predict the RAMPAGE or CAGE support in another cell type, in a cross-cell type manner (Fig. S6f). This approach may be used to define a set of high-confidence TSS regions from LR-RNA-seq that would also be supported by RAMPAGE or CAGE where neither RAMPAGE nor CAGE data are available in the cell type of interest. This demonstrates that TSSs derived from LR-RNA-seq serve as a reasonable stand-in for CAGE and RAMPAGE, with the added benefit that LR-RNA-seq profiles both ends and the exon structure of transcripts at the same time.

### Applying Cerberus to the human ENCODE4 LR-RNA-seq dataset leads to the largest number of detected alternative splicing events to date

We compared the detection of alternative splicing (AS) events in our dataset with a recent LR-RNA-seq transcriptome published by the GTEx consortium ^23^. We ran SUPPA2^43^ on the observed LR-RNA-seq transcripts and obtained, for every gene and type of local AS event, a list of AS transcripts. We found a considerably larger number of AS transcripts compared to those reported in the GTEx LR-RNA-seq catalog. We observed a higher proportion of novel AS transcripts defined by EC compared to TSS and TES (Fig. S8a), albeit lower than those reported by GTEx. This is likely due to the fact that our novel transcripts are defined with respect to a more recent GENCODE version (v40) than the one used by the GTEx study (v26). In support of this, we found that the majority of our observed ENCODE LR-RNA-seq transcripts, both known and novel, are missing in the GTEx catalog (Fig. S8b). On the other hand, although most of the GTEx novel transcripts are not reported in the ENCODE4 catalog, they represent combinations of already annotated splice junctions (NIC). From a methodological perspective, we also found that Cerberus accounts for a larger variety of AS events related to TSSs and TESs (0.25 < PSI < 0.75) compared to SUPPA2 (Fig. S8c). Altogether, this indicates that the ENCODE4 LR-RNA-seq catalog provides the largest set of novel and annotated AS events in the human transcriptome available to date.

## SUPPLEMENTARY METHODS

### B6/Cast mouse tissue collection

Mouse tissues were harvested from C57BL/6J (RRID:IMSR_JAX:000664) x CAST/EiJ (RRID:IMSR_JAX:000928) F1 animals across 7 postnatal day (PND) or postnatal month (PNM) timepoints: PND4, PND10, PND14, PND25, PND36, 2 months and 18-20 months. Tissues were flash frozen in liquid nitrogen and stored at -80C prior to processing.

### Preprocessing short-read RNA-seq data and data availability

All short-read RNA-seq data was preprocessed according to the details on the ENCODE portal. Gene quantification of 548 short RNA-seq datasets were downloaded from the ENCODE portal using this cart (https://www.encodeproject.org/carts/4ea7a43f-e564-4656-a0de-b09c92215e52/), then TPM values for polyA genes were extracted from each of them.

### Preprocessing microRNA-seq data and data availability

Quantification of 254 microRNA-seq datasets using GENCODE GRCh38 V29 annotations were downloaded from the ENCODE portal using this cart (https://www.encodeproject.org/carts/human_mirna/). Counts were concatenated across all datasets and converted to CPM for downstream analyses.

### Preprocessing LR-RNA-seq data and data availability

All LR-RNA-seq data was preprocessed according to the details on the ENCODE portal. Input and output files, including the final Cerberus GTFs, gene triplets, and transcript triplets, are available at the following accessions:

- Human: ENCSR957LMA
- Mouse: ENCSR110KDI

Raw data are available at the following links:

- Human: https://www.encodeproject.org/carts/829d339c-913c-4773-8001-80130796a367/
- Mouse: https://www.encodeproject.org/carts/55367842-f225-45cf-bfbe-5ba5e4182768/

### Human / Mouse LR-RNA-seq annotation with TALON and LAPA

Mapped LR-RNA-seq BAMs were obtained from the ENCODE portal using the above cart links for human and mouse respectively. Reads were annotated with their 3’ end A content using the `talon_label_reads` module and hg38 / mm10. Reads were annotated using talon with reference annotation GENCODE v29 / vM21. Output transcripts were filtered for reproducibility of 5 reads across 2 libraries, and for reads that had fewer than 50% A nucleotides in the last 20 bp of the 3’ end to remove artifacts of internal priming using the `talon_filter_transcripts` command. Unfiltered and filtered transcript abundance matrices were obtained using the `talon_abundance` command. A filtered GTF was obtained using the `talon_create_GTF` command. From the unfiltered TALON abundance, counts of each gene were computed by summing up counts for each transcript per gene.

We ran LAPA on the bam files mentioned above to create TSS and TES clusters from LR-RNA-seq. If the bam files had replicates, we filtered clusters by choosing a cutoff that ensures a 95% replication rate. Samples without replicates were filtered with a median cutoff of replicated clusters. Using those TSS and TES clusters and the read_annot created by TALON, we corrected TSSs and TESs of the filtered TALON GTF file. During the correction, new transcript isoforms were created if the same exon junction chain mapped to multiple start and end sites.

### Gene rank analysis

For detected (>= 1 TPM in any library) polyA genes in the human LR-RNA-seq dataset, we ranked the genes in each library according to their expression (1 = most highly expressed) and plotted the genes at specific ranks for each library by their TPM, split by cell line and tissue derived libraries. For statistical testing between the cell line and tissue groups, we performed a Wilcoxon rank-sum test with p-value thresholds P > 0.05; *P <= 0.05, **P <= 0.01, ***P <= 0.001, ****P <= 0.0001.

### Novel gene analysis

For novel genes in both human and mouse, we first filtered our novel TALON transcripts for those that passed the filters previously described (5 reads in at least 2 libraries and <50% A nucleotides in the last 20bp of the 3’ end). We then selected only the transcripts that passed this filter that belonged to novel intergenic genes and that had at least one spliced (i.e. more than one exon) transcript isoform expressed >= 1 TPM. To make an analogous comparison to our annotated genes, we performed the same filtering on our TALON transcripts with the exception of requiring the transcripts to be from annotated polyA genes rather than from novel intergenic genes.

### Cerberus overview

#### Obtaining annotated TSS / TES regions from GTFs

Given a GTF, `cerberus gtf_to_bed` (Fig. S4a) will extract the single base pair TSS and TES coordinates and extend them by n bp on either side. Regions within m bp of one another are merged. Each unique combination of coordinates, strand, and gene is recorded in BED format.

#### Obtaining annotated exon junction chains from GTFs

Given a GTF, `cerberus gtf_to_ics` will extract each unique combination of intron coordinates, strand, and gene and record them in a tab-separated format (Fig. S4a).

#### Assigning triplet features numbers

As part of both `cerberus gtf_to_ends` and `gtf_to_ics` (Fig. S4a), Cerberus numbers triplet features based on their annotation status within the reference GTF, if any. For triplet features derived from these GTFs, each TSS, EC, and TES is numbered from 1 to n within each gene based on the annotation status of the transcript they were derived from. Transcripts are first ordered by MANE status, then APPRIS ^44^ principal status, and finally whether the transcript comes from the GENCODE basic set. The result is that triplet features from MANE transcripts are always numbered 1, and lower triplet feature numbers within a gene correspond to transcripts with more importance as determined by GENCODE.

#### Merging TSS and TES regions across multiple BED files

For each BED file input `cerberus agg_ends` (Fig. S4a) takes a boolean argument for whether the regions should be used to initialize new TSS / TES regions, a boolean argument for whether the regions should be considered reference regions, and a name for each BED file source. BED files without a gene identifier cannot be used to initialize regions. For the first BED file, Cerberus creates a set of reference regions and uses the triplet feature numbers that were previously assigned by Cerberus to name each TSS or TES. The first BED file must have gene IDs and must be used to initialize the regions. For each subsequent BED file, in order, Cerberus determines which new regions are within m bp of a region already in the reference. These regions are added as sources of support for the already-existing regions, but do not extend the boundaries of existing regions in order to combat growing regions as more data is added. If a region is not within m bp of an existing region and the initialize regions option is turned on, the new region is added as a new region in the reference set. After all new regions have been added, triplet feature numbers are computed by ordering the features within each gene based on the number assigned by Cerberus in a previous step and then incrementing the preexisting Cerberus reference maximum number. BED files that are not used to initialize new regions will only ever be added as additional forms of support for each region already in the reference.

#### Merging ECs across multiple EC files

For each EC file, `cerberus agg_ics` (Fig. S4a) takes a boolean argument for whether the ECs should be used as a reference and a source name. For the first EC file, Cerberus creates a reference set of ECs and uses the EC numbers that were determined using `cerberus gtf_to_ics`. For each subsequent EC file, Cerberus finds ECs that are not already in the Cerberus reference set, orders the new ECs by their numbers from `cerberus gtf_to_ics`, creates new numbers for each EC by, in order, assigning them numbers by incrementing from the maximum existing number for a gene from the reference.

#### Creating a Cerberus reference

After generating an aggregated TSS, EC, and TES file, `cerberus write_reference` (Fig. S4a) will write all three tables in a Cerberus reference h5 format, a well-supported and commonly used data structure that can store multiple tables.

#### Updating a GTFs and counts matrices with a Cerberus annotation

After each transcript from a transcriptome has been assigned a transcript triplet, the corresponding GTF and counts matrix from the transcriptome can be updated to use the new transcript identifier using `cerberus replace_gtf_ids` and `cerberus replace_ab_ids` (Fig. S4b). Cerberus will replace the transcript ids with the transcript triplets and, if requested, merge transcripts that are assigned duplicate transcript triplets, summing the counts in the case of the counts matrix.

#### Gene triplet and gene structure simplex coordinate computations

Following transcriptome annotation, gene triplets can be calculated for different sets of annotated transcripts using Cerberus’ Python API and the `CerberusAnnotation data structure`. Regardless of the input set, Cerberus computes the gene triplets by counting the number of unique TSSs, ECs, and TESs used across a set of transcripts (Fig. 3a, Fig. S9). This calculation can be done without any filtering using the `CerberusAnnotation.get_source_triplets()` function, which computes the number of TSSs, ECs, and TESs used across each transcriptome annotated by Cerberus. `CerberusAnnotation.get_expressed_triplets()` will calculate the gene triplets for individual samples based on the subset of transcripts that are expressed in each sample and can optionally use a table of transcript / sample combinations to determine which transcripts are used in each sample. Finally, `CerberusAnnotation.get_subset_triplets()` simply takes in a list of transcripts to compute a gene triplet for the entire input set. In all cases, the number of transcripts used to calculate the gene triplet is also recorded. After computing the gene triplets, the EC count is converted to the splicing ratio. To generate the gene structure simplex coordinates, the sum of the number of TSSs, splicing ratio, and number of TESs is normalized such that they sum to one (Fig. 3a-b, Fig. S9).

Additionally, the sector assignments are generated for each gene triplet. Genes with a TSS simplex coordinate >0.5 are TSS-high, those with a TES simplex coordinate >0.5 are TES-high, and those with a splicing ratio simplex coordinate >0.5 are splicing-high.Genes where all three simplex coordinates <= 0.5 are mixed, and genes with just one transcript are in the simple sector. An important note is that mixed genes can have the same coordinates as a simple gene. To this end, when calculating gene triplets, the number of transcripts used to generate the triplet is also recorded and used to separate out the simple from the mixed genes (Fig. 3a-b, Fig. S9).

#### Computing gene triplet centroids

Given a set of gene triplets, the centroid is computed by averaging each gene structure simplex coordinate. The resulting coordinate retains the property that it sums to one (Fig. S9).

#### Computing distances in the gene structure simplex

We compute the distance between any two points on the gene structure simplex as the Jensen-Shannon distance (Fig. S9). Jensen-Shannon distance is a metric on probability distributions ^45^. For a given pair of gene structure simplex coordinates, the Jensen-Shannon distance is computed in Cerberus using Scipy ^46^ with the `scipy.spatial.distance.jensenshannon` function.

### Cerberus processing of human ENCODE4 LR-RNA-seq dataset

#### Obtaining annotated TSS / TES regions from GTFs

The GTF files from GENCODE v40, GENCODE v29, the LAPA output GTF representing the human ENCODE LR-RNA-seq dataset, and the GTEx LR-RNA-seq GTF were used to obtain TSS and TES regions associated with each transcript using `cerberus gtf_to_ends`. For each GTF, the single base pair TSS and TES coordinates were extracted and extended 50 bp on either side, and regions within 50 bp of one another were merged. Each unique combination of coordinates, strand, and gene were recorded.

#### Obtaining external TSS / TES regions

External datasets used to support TSSs were obtained from the ENCODE CAGE and RAMPAGE data, FANTOM CAGE data4, and ENCODE PLS, pELS, and dELS cCREs. External datasets used to support TESs were obtained from ENCODE PAS-seq data, and the PolyA Atlas ^18^. Each file was downloaded in BED format and converted to the BED format required for Cerberus.

#### Obtaining annotated exon junction chains from GTFs

The GTF files from GENCODE v40, GENCODE v29, from the human ENCODE LR-RNA-seq output GTF, and the GTEx LR-RNA-seq GTF were used to obtain exon junction chains from each transcript using `cerberus gtf_to_ics`. Each unique combination of intron coordinates, strand, and gene were recorded.

#### Creating a set of reference triplet features

To create a consensus reference set of triplet features, `cerberus agg_ends` and `cerberus agg_ics` (Fig. S4a) were run on the aforementioned TSS, EC, and TES sets, with m=20 for the TSSs and TESs. The triplet features from GENCODE v40 and v29 were used as reference features. For the TSSs, new regions were incorporated from GENCODE v40, v29, the human ENCODE LR-RNA-seq data, and the GTEx data, whereas the CAGE, RAMPAGE, and cCRE data were only used as forms of support for existing regions. For TESs, new regions were incorporated from GENCODE v40, v29, the human ENCODE LR-RNA-seq data, and the GTEx data, whereas the PAS-seq and PolyA Atlas regions were used as forms of support for existing regions.

#### Transcriptome annotation

The GTFs of the GENCODE v40, GENCODE v29, and human ENCODE LR-RNA-seq transcriptomes were annotated with `cerberus annotate_transciptome`, updated GTFs were generated with `cerberus replace_gtf_ids` with the update ends and collapse options used (Fig. S4b). For the human ENCODE LR-RNA-seq data, `cerberus replace_ab_ids` was also run on the filtered abundance file output from LAPA using the collapse option to generate a matching counts matrix (Fig. S4b).

### Cerberus analysis of human ENCODE4 LR-RNA-seq

#### Finding observed transcripts and transcripts expressed in a sample

Observed transcripts are defined as transcripts that are expressed >= 1 TPM in any given library. Observed transcripts in a specific sample are transcripts that are expressed >= 1 TPM in any library that belongs to the same sample.

#### Finding observed major transcripts and major transcripts in a sample

For each sample, each transcript is assigned a percent isoform (pi, 0-100) value that indicates what percentage of the gene’s expression is derived from said transcript using Swan ^47^. Transcripts for a gene are then ranked by pi value. In order from the highest pi value transcript to the lowest pi value transcript, transcripts are added to the major transcript set until the cumulative pi value of the set is >90, yielding the samplelevel major transcript set. The observed major transcripts for the entire dataset is computed by taking the union of all major transcripts across all samples. In both cases, transcripts are limited to those that have passed the observed and sample-level observed transcripts as defined above.

#### Gene triplet computations

Gene triplets were calculated for the following sets of transcripts, all just using polyA genes:

- All transcripts from annotated GENCODE v40 genes (v40)
- All observed transcripts (observed)
- All observed major transcripts (observed major)
- Detected transcripts in each sample (sample-level)
- Detected major transcripts in each sample (sample-level major)
- All observed transcripts in the dataset from samples that match the mouse samples (mouse match)
- All observed major transcripts in the dataset from samples that match the mouse samples (mouse match major)

#### Transcriptional diversity by gene biotype comparison

Using the gene triplets table, we found the gene / sample combination where each polyA gene is most highly expressed and recorded the gene TPM and number of transcripts from that gene in that sample. We then split each gene into its biotype category (protein coding, lncRNA, or pseudogene) and into a TPM bin (lowly expressed, 1-10 TPM; medium expressed, 10-100 TPM; and highly expressed, 100-max TPM).

#### Gene structure simplex distances computed

We computed the follow pairwise distances between simplex points:

- Sample-level gene triplet vs. the centroid for the sample-level gene triplets
- Observed gene triplet vs. the centroid for the sample-level gene triplets for each gene with at least 2 transcripts
- Observed gene triplet vs. observed major gene triplet

Each set of distances was computed using only protein coding genes. Z-scores were also computed for each comparison.

#### Comparing sample-level to observed gene triplets

The number of transcripts, TSSs, ECs, and TESs was calculated for each gene globally (i.e. transcripts or triplet features / gene) and for each sample (i.e. transcripts or triplet features / gene / sample). For transcripts, TSSs, ECs, and TESs separately, a two-sided KS test was performed using Scipy’s `stats.kstest` function to assess statistical differences between the global and sample-level transcripts or triplet features per gene distributions.

#### Calling predominant transcripts

On both the sample and library level, we called the most highly expressed transcript from a gene the predominant transcript for that gene. On the sample level, we used the mean expression of the transcript.

#### Predominant transcript MANE comparison

We first restricted this analysis to only consider genes which have annotated MANE transcripts. For these genes, we determined how often the predominant transcript for a given gene is the MANE transcript for a gene in each library.

### Cerberus processing of mouse ENCODE4 LR-RNA-seq dataset

#### Obtaining annotated TSS / TES regions from GTFs

The GTF files from GENCODE vM25, GENCODE vM21, and from the LAPA output GTF representing the mouse ENCODE LR-RNA-seq dataset were used to obtain TSS and TES regions associated with each transcript using `cerberus gtf_to_ends`. For each GTF, the single base pair TSS and TES coordinates were extracted and extended 50 bp on either side, and regions within 50 bp of one another were merged. Each unique combination of coordinates, strand, and gene were recorded.

#### Obtaining external TSS / TES regions

External datasets used to support TSSs were obtained from the ENCODE mouse PLS, pELS, and dELS cCREs. External datasets used to support TESs were obtained from the mouse PolyA Atlas. Each file was downloaded in BED format and converted to the BED format required for Cerberus.

#### Obtaining annotated exon junction chains from GTFs

The GTF files from GENCODE vM25, GENCODE vM21, and from the mouse LR-RNA-seq GTF were used to obtain exon junction chains from each transcript using `cerberus gtf_-to_ics`. Each unique combination of intron coordinates, strand, and gene were recorded.

#### Creating a set of reference triplet features

To create a consensus reference set of triplet features, `cerberus agg_ends` and `cerberus agg_ics` (Fig. S4a) were run on the aforementioned TSS, EC, and TES sets, with m=20 for the TSSs and TESs. The triplet features from GENCODE vM25 and vM21 were used as reference features. New TSSs were incorporated from GENCODE vM25, vM21, the mouse ENCODE LR-RNA-seq data, whereas the cCRE data were only used as forms of support for existing regions. New TESs were incorporated from GENCODE vM25, vM21, and the mouse ENCODE LR-RNA-seq data, whereas the PolyA Atlas regions were used as forms of support for existing regions.

#### Transcriptome annotation

The GTFs of the GENCODE vM25, GENCODE vM21, and mouse ENCODE LR-RNA-seq transcriptomes were annotated with `cerberus annotate_transciptome`, updated GTFs were generated with `cerberus replace_gtf_ids` with the update ends and collapse options used (Fig. S4b). For the mouse ENCODE LR-RNA-seq data, `cerberus replace_ab_ids` was also run on the filtered abundance file output from LAPA using the collapse option to generate a matching counts matrix (Fig. S4b).

### Cerberus analysis of mouse ENCODE4 LR-RNA-seq

#### Finding observed transcripts and transcripts expressed in a sample

Observed transcripts are defined as transcripts that are expressed >= 1 TPM in any given library. Observed transcripts in a specific sample are transcripts that are expressed >= 1 TPM in any library that belongs to the same sample.

#### Finding observed major transcripts and major transcripts in a sample

For each sample, each transcript is assigned a percent isoform (pi, 0-100) value that indicates what percentage of the gene’s expression is derived from said transcript using Swan ^47^. Transcripts for a gene are then ranked by pi value. In order from the highest pi value transcript to the lowest pi value transcript, transcripts are added to the major transcript set until the cumulative pi value of the set is >90, yielding the samplelevel major transcript set. The observed major transcripts for the entire dataset is computed by taking the union of all major transcripts across all samples. In both cases, transcripts are limited to those that have passed the observed and sample-level observed transcripts as defined above.

#### Gene triplet computations

Gene triplets were calculated for the following sets of transcripts; all just using polyA genes:

- All annotated GENCODE vM25 genes (vM25)
- All observed transcripts in the dataset (observed)
- All observed major transcripts in the dataset (observed major)
- Detected transcripts in each sample (sample-level)
- Detected major transcripts in each sample (sample-level major)

#### Calling predominant transcripts

On both the sample and library level, we called the most highly expressed transcript from a gene the predominant transcript for that gene. On the sample level, we used the mean expression of the transcript.

#### Human-mouse comparison

We found orthologous genes between human and mouse using this Biomart query, and subset our considered genes to those that were protein coding, expressed in both species, and were just 1:1 orthologs. We determined the sector of each gene in each species using the observed major gene triplets in mouse, and the mouse match major gene triplets in human. We counted the number of genes that have the same sector between human and mouse. Furthermore, we compared the sector of each orthologous pair of genes between species just in the matching embryonic stem cell samples (H1 in human, F121-9 in mouse) between human and mouse to verify that the trend seen overall was reproducible on a more one to one comparison. Additionally, we computed the centroids from the sample-level gene triplets from matching samples in human and all sample-level gene triplets in mouse mouse and calculated the Jensen-Shannon distances between sample-level centroids for each orthologous gene in human and mouse.

#### ORF and NMD prediction

We used TAMA’s ^48^ ORF / NMD prediction pipeline with [minimal changes](https://github.com/fairliereese/tama) to support our file formats. To pick one representative ORF from each transcript, we chose the ORF with the highest percent identity from BLASTP ^49^ to an annotated GENCODE v40 protein sequence; breaking ties by considering ORF completeness. For transcripts with no BLASTP hits to known transcripts, we picked complete ORFs; breaking ties by picking the longest ORF.

#### Comparing detection of AS events by SUPPA and Cerberus

We used SUPPA2 (v2.3^43^) to define alternative splicing (AS) events (A3: alternative 3’ splicing; A5: alternative 5’ splicing; AF: first exon; AL: last exon; IR: intron retention; SE: exon skipping; MX: mutually exclusive exons). Specifically, we generated a catalog of local AS events based on the Cerberus GTF file (function `generateEvents`) and used the novelties of each observed transcript to compute the proportion of novel transcripts (based on EC, TSS, or TES) out of the total set of transcripts involved in a particular type of event.

Next, we used SUPPA2 to compute the Proportion of Splicing Index (PSI) for each type of event using the observed transcript filtered expression matrix (polyA transcripts expressed >=1 TPM in at least one library; function `psiPerEvent`). PSI values were averaged between replicates of the same sample. We selected genes with at least one local AF or AL event, applying a threshold of 0.25 < PSI < 0.75.

In order to compare the detection of AS events by SUPPA2 and Cerberus, we also computed triplet feature PSI values based on events identified by Cerberus by dividing the counts for any given TSS, EC, or TES by the total counts for the gene in a given sample. We selected genes with at least one local event at the TSS or TES (0.25 < PSI < 0.75). Next, we computed the intersection between genes showing AF (SUPPA2) and TSS (Cerberus) events, and between genes showing AL (SUPPA2) and TES (Cerberus) events.

#### Machine learning models for RAMPAGE and CAGE TSS prediction

RAMPAGE and CAGE TSS annotation data for GM12878 and K562 were obtained from ENCODE portal (ENCSR000AEI, ENCSR000AER, ENCSR000CJN, ENCSR000CKA). LAPA and Cerberus TSS regions derived from just one experiment each for GM12878 and K562 (ENCSR962BVU and ENCSR589FUJ respectively) were used for long-read data. Using bedtools intersect ^50^ a binary (0/1) label for each long-read peak was assigned depending on whether the region overlapped with at least one peak in either of the RAMPAGE or CAGE assays in the same cell type. Average DHS signal values over LR TSS peaks were calculated using UCSC `bigWigAverageOverBed` on GM12878 and K562 DHS-seq experiments (ENCSR000EMT, ENCSR000EOT). Test sets include long-read regions from chromosomes 2 and 3, whereas training sets include all other human chromosomes. 7 logistic regression models were trained on each long-read experiment using all the 2^3^-1 combinations of peak’s TPM expression, DHS signal, and length (i.e. in R: glm(label TPM + DHS + length, type =“binomial”)) where the input parameters have been log2-transformed) and the AIC values were calculated and ranked in each experiment and for each model type. The model using all 3 parameters (TPM, DHS, and length) had the lowest AIC, meaning that given the number of parameters and the observed RSS error, logit[label TPM + DHS + length] had the highest predictive power and was therefore selected. For the same cell-type prediction, a model is trained on [chr1, chr4-22, chrX] and tested on [chr2, chr3] for long-read data from the same cell type (ex: K562). In cross-cell type prediction, a model is trained and tested on two different cell lines (ex: trained on [chr1, chr4-22, chrX] of a GM12878 long-read experiment and tested on [chr2, chr3] of a K562 long-read experiment). Cerberus replicates belonging to the same experiment were combined by taking the average mean-normalized TPM values of the identical peaks across different replicates.

## Data and code availability

- Human LR-RNA-seq data / processing pipeline: https://www.encodeproject.org/annotations/ENCSR957LMA/
- Mouse LR-RNA-seq data / processing pipeline: https://www.encodeproject.org/annotations/

ENCSR110KDI

- Processing / figure generation code: https://github.com/fairliereese/paper_rnawg
- Cerberus: https://github.com/fairliereese/cerberus

## SUPPLEMENTARY TABLES

- Table S1: Human LR-RNA-seq library metadata.
- Table S2: Mouse LR-RNA-seq library metadata.

## SUPPLEMENTARY FIGURES

**Figure S1.**
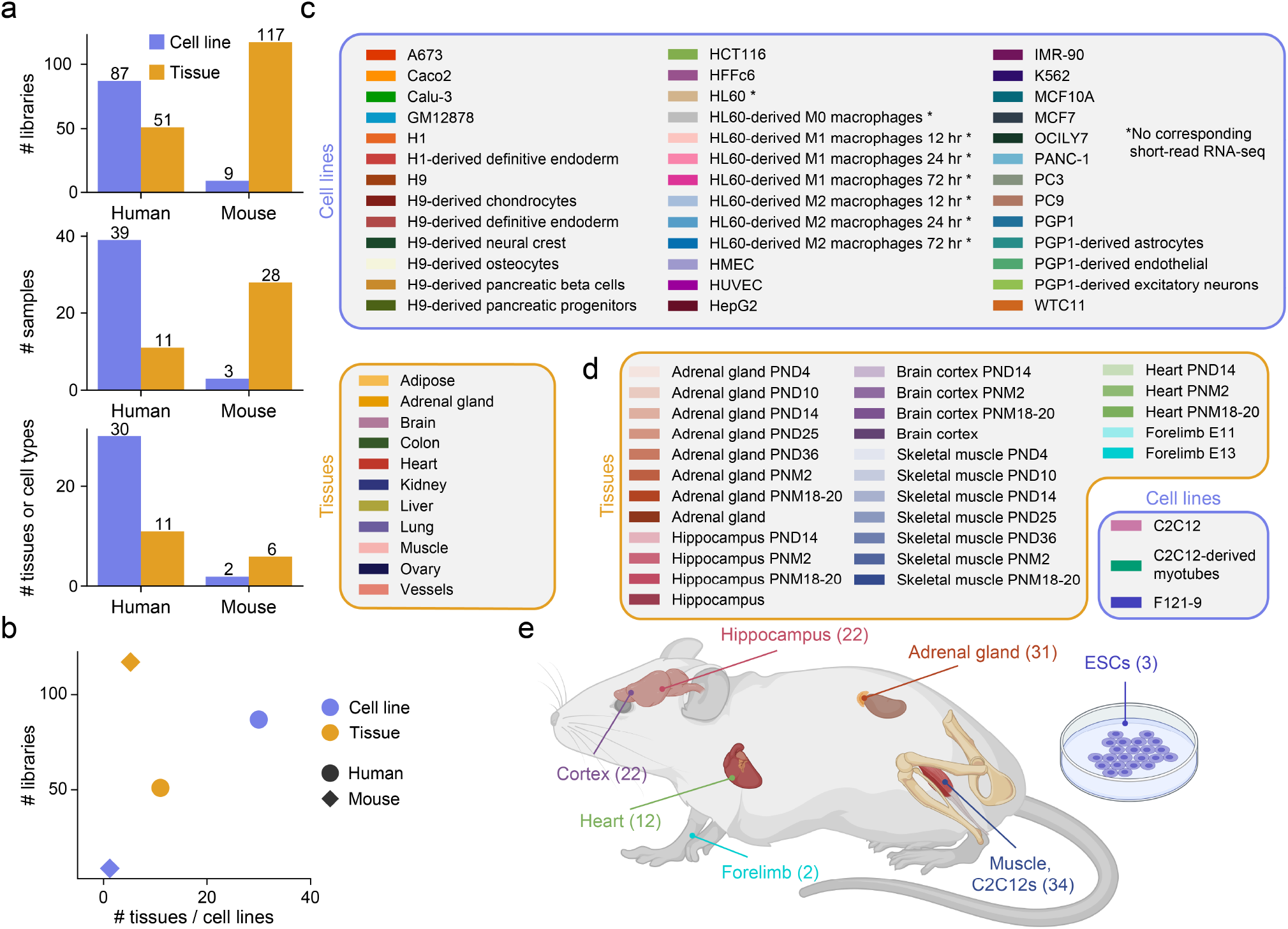
Overview of the ENCODE4 LR-RNA-seq dataset. a, From top to bottom, number of LR-RNA-seq libraries, samples (split by cell line / tissue identity as well as timepoint, when relevant), and unique tissues or cell types in the ENCODE LR-RNA-seq dataset split by species and tissue or cell line. b, Number of LR-RNA-seq libraries versus the number of tissues or cell lines assayed, split by species and cell line / tissue. c-d, Color legend and labels for each c, human sample, with samples that lack corresponding short-read RNA-seq data denoted by a star d, mouse sample in the LR-RNA-seq dataset; split by tissues and cell lines. e, Overview of the sampled tissues and number of libraries from each tissue in the ENCODE mouse LR-RNA-seq dataset.

**Figure S2.**
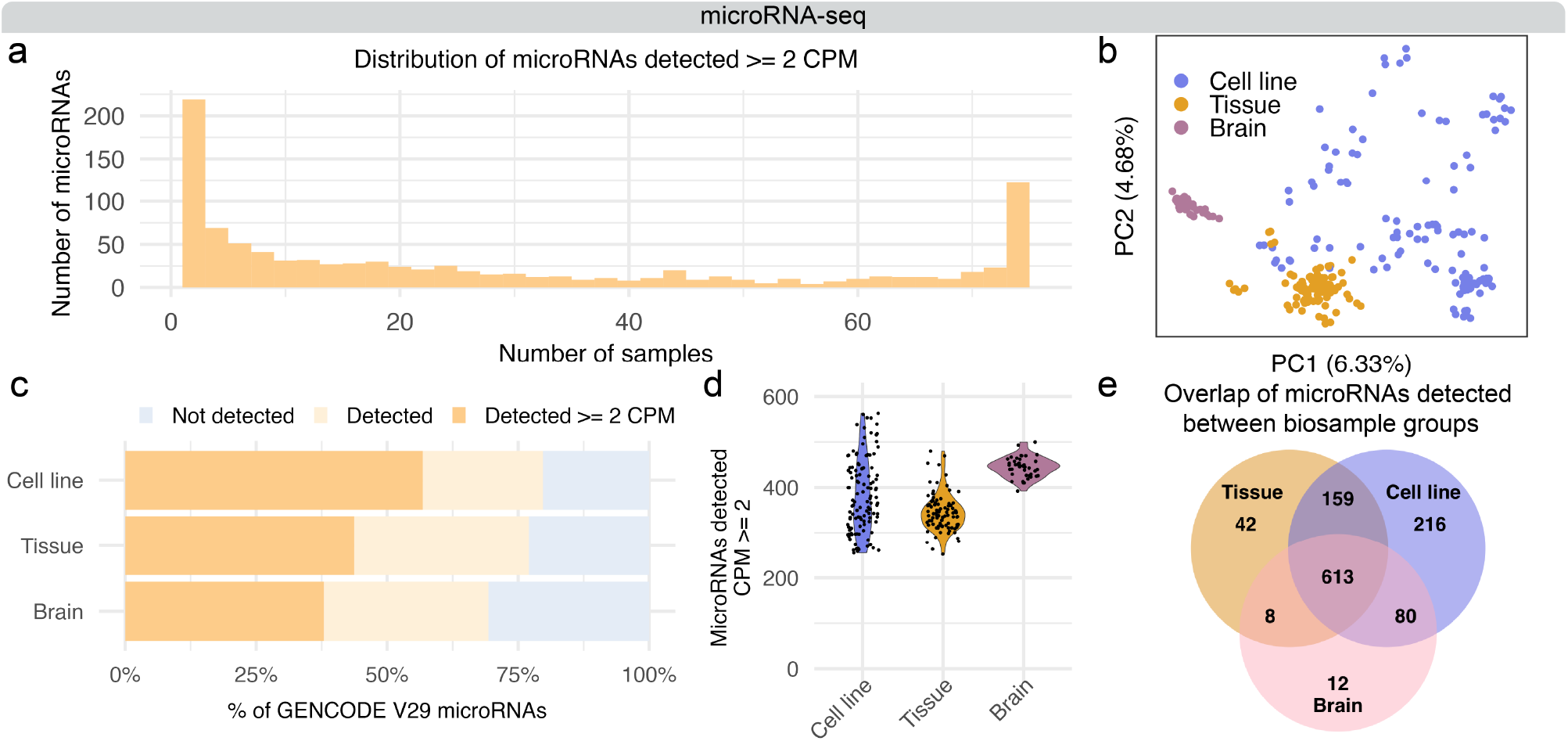
Overview and detection of microRNAs in the ENCODE microRNA-seq dataset. a, Distribution of GENCODE v29 mature microRNAs detected at CPM > 2 between cell lines, tissues, and brain tissue samples. b, PCA computed on microRNAs detected >2 CPM in each human microRNA-seq library, colored by cell line and tissue designation and by brain tissue. c, Percentage of GENCODE v29 microRNAs detected in at least one ENCODE human microRNA-seq library from either cell line, tissue, or brain tissue samples at > 0 CPM and > 2 CPM. d, Number of samples in which each GENCODE v29 microRNA is detected at > 2 CPM in the ENCODE human microRNA-seq dataset. e, Overlap of detected (> 2 CPM) microRNAs in at least one library derived from cell line, tissue, or brain tissue.

**Figure S3.**
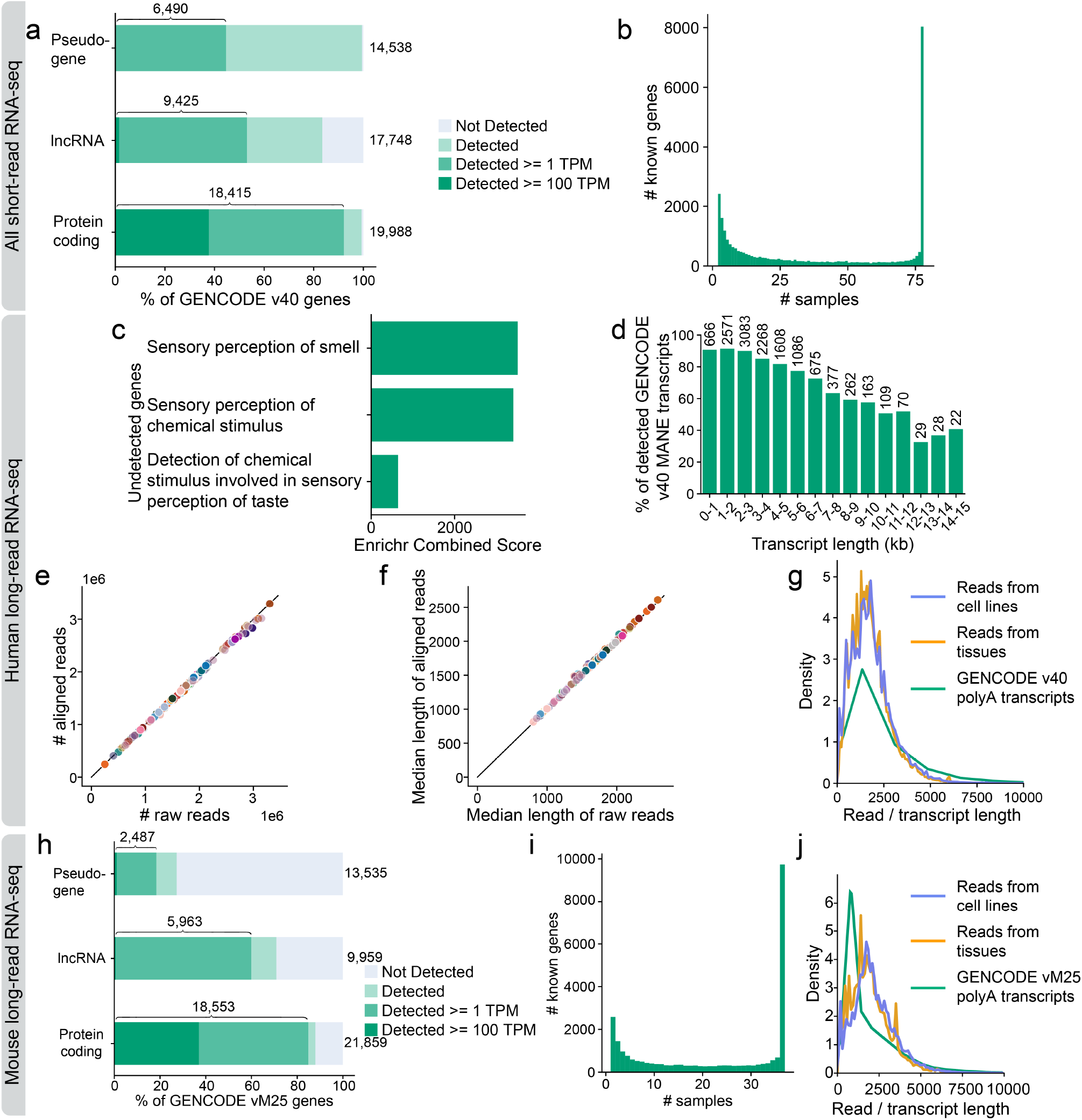
Gene detection from short-read RNA-seq; gene detection, read length and alignment QC in both human and mouse LR-RNA-seq. a, Percentage of GENCODE v40 polyA genes by gene biotype detected in at least one ENCODE short-read RNA-seq library from all samples at >0 TPM, >=1 TPM, and >=100 TPM. b, Number of samples in which each GENCODE v40 gene is detected >= 1 TPM in the ENCODE short-read RNA-seq dataset in all samples. c, Top 3 biological process GO terms from GENCODE v40 protein coding genes that were not detected in the human LR-RNA-seq dataset. d, Percentage (y-axis) and number (top of each bar) of GENCODE v40 MANE transcripts we detect binned by transcript length. Restricted to expressed genes that we detect >= 10 TPM in at least one library. e, Number of raw reads vs. number of aligned reads in each human LR-RNA-seq library. f, Median length of each raw read vs. median length of the aligned portion of each read in each human LR-RNA-seq library. g, Read length profiles of post-TALON reads from LR-RNA-seq data split by tissue or cell line designation and polyA transcript length profile from GENCODE v40. h, Percentage of GENCODE vM25 polyA genes by gene biotype detected in at least one ENCODE mouse LR-RNA-seq library at >0 TPM, >=1 TPM, and >=100 TPM. i, Number of samples that each GENCODE vM25 gene is detected >= 1 TPM in the ENCODE mouse LR-RNA-seq dataset. j, Read length profiles of post-TALON reads from mouse LR-RNA-seq data split by tissue or cell line designation and polyA transcript length profile from GENCODE vM25.

**Figure S4.**
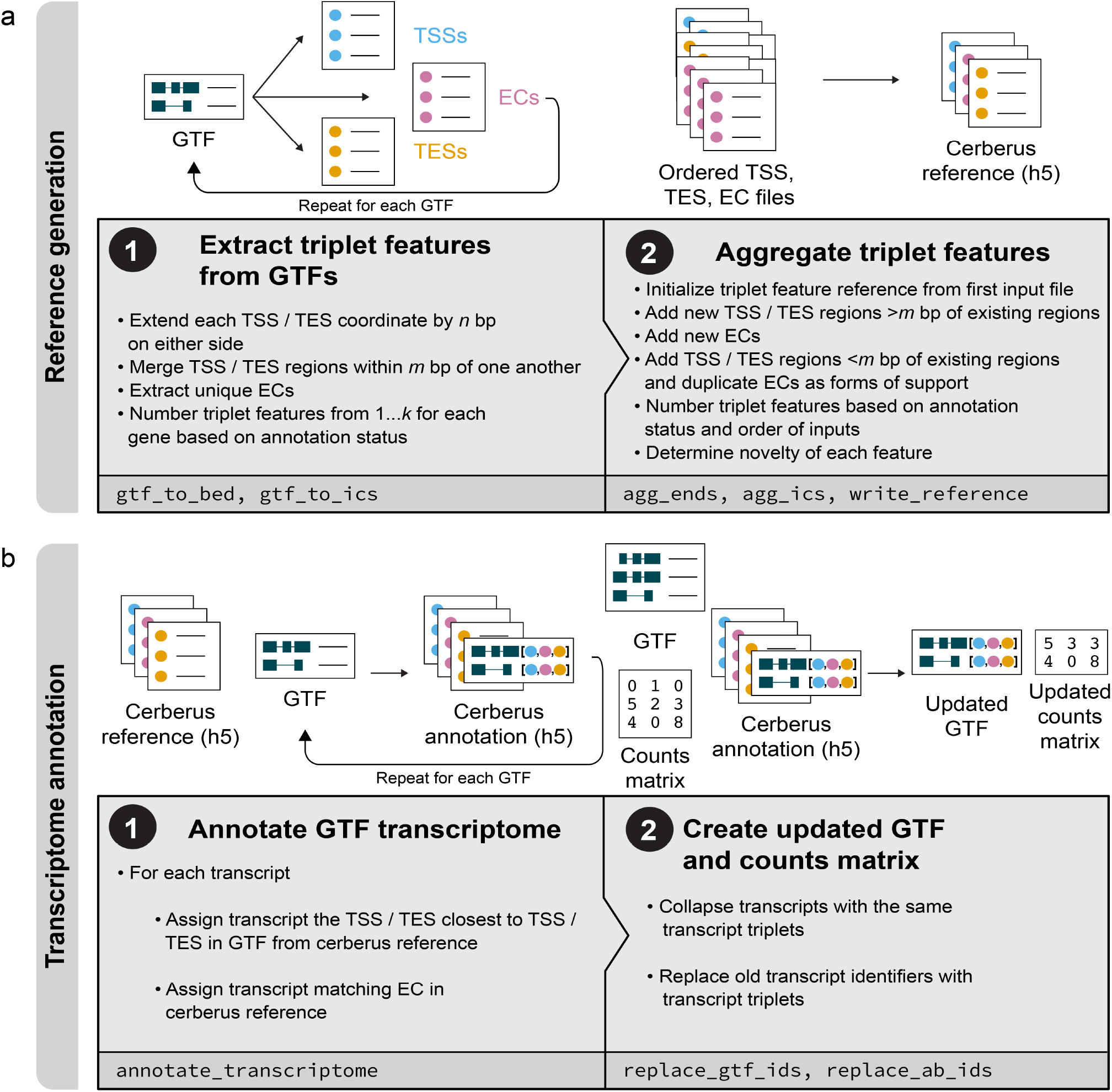
Overview of Cerberus processing of transcriptomes and triplet features. a, Workflow for generating a Cerberus reference: a collection of TSSs, ECs, and TESs (triplet features) sourced from various inputs. b, Workflow for generating a Cerberus transcriptome annotation, which assigns each transcript in a GTF a set of triplet features (TSS, EC, TES) from the Cerberus reference.

**Figure S5.**
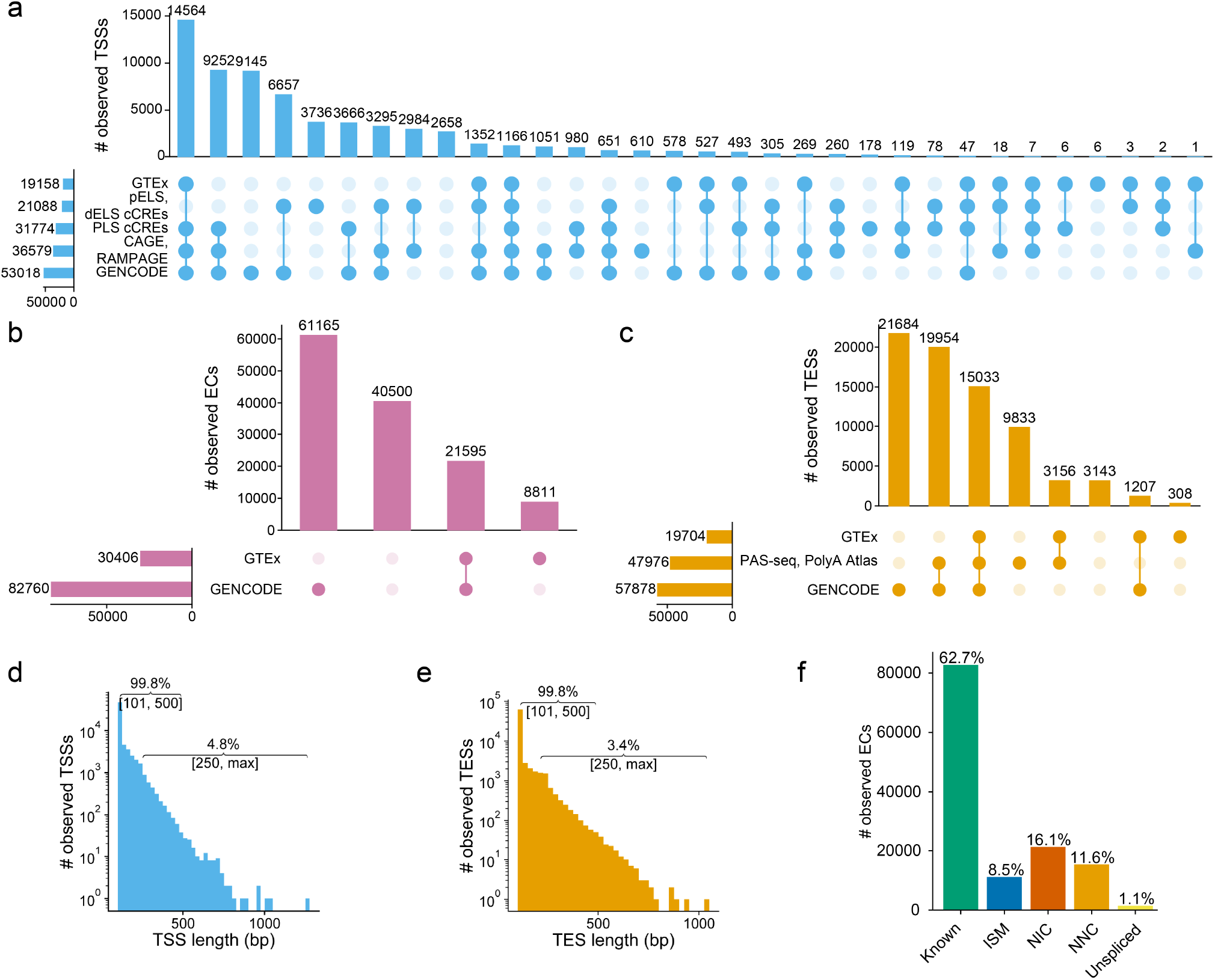
Characterization of observed triplet features from human LR-RNA-seq. a-c,Upset plots showing sources that overlap observed triplet features derived from human ENCODE LR-RNA-seq for a, TSSs b, ECs c, TESs. d-e, Lengths of observed d, TSSs e, TESs derived from human ENCODE LR-RNA-seq. f Novelty of unique ECs detected >= 1 TPM from polyA genes in human ENCODE LR-RNA-seq.

**Figure S6.**
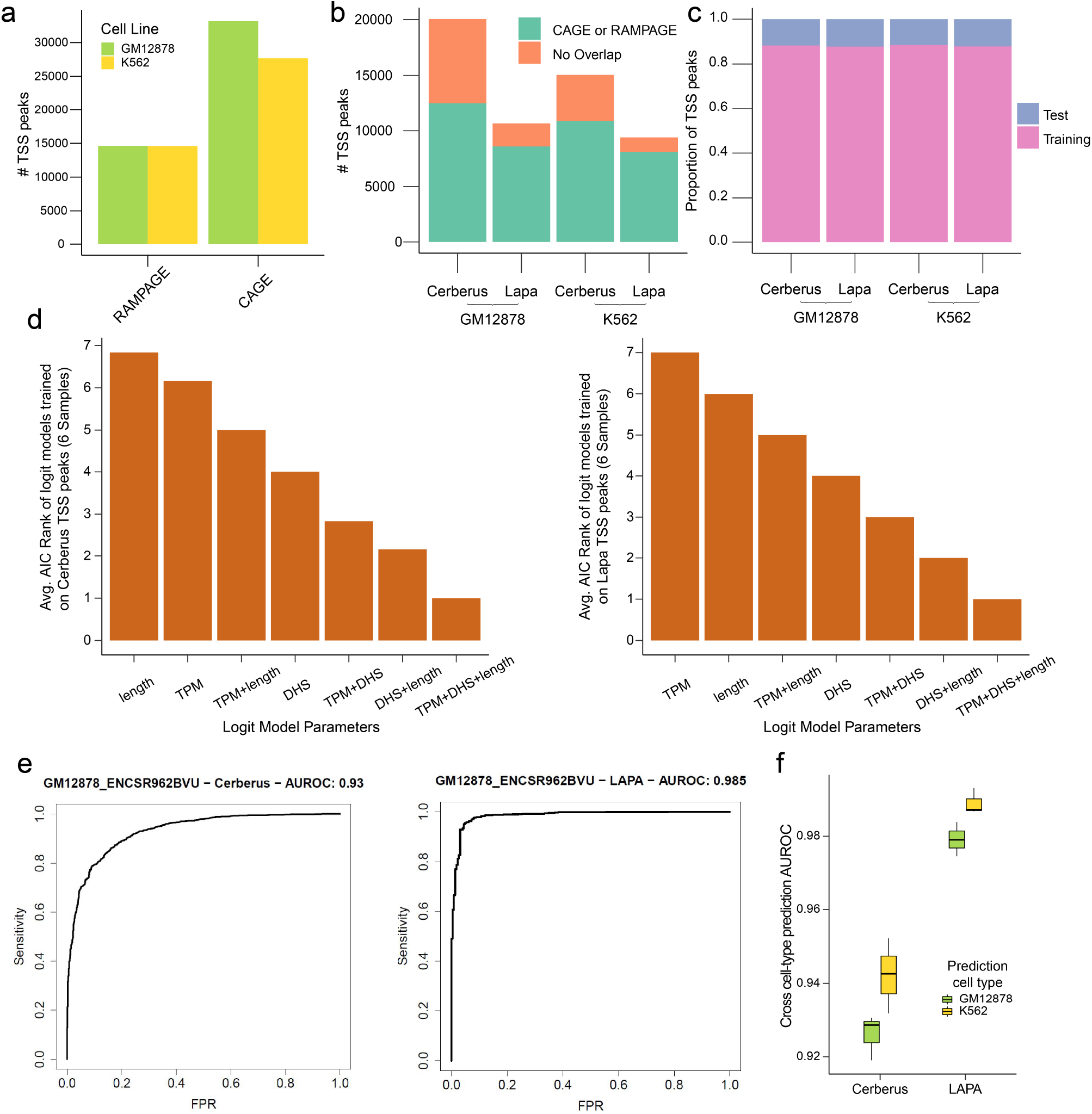
Machine learning models predict support for long-read TSS peaks by other TSS assays in a cross-cell type manner. a, Number of long-read RNA-seq TSS peaks called by RAMPAGE and CAGE. b, Number of long-read TSS peaks called by Cerberus or LAPA supported by RAMPAGE or CAGE assays in GM12878 and K562 long-read experiments gm12878_3 and k562_2. c, Fraction of peaks used for the test (chr 2 and chr3) and training sets (all other chromosomes) in K562 and GM12878 long-read experiments. d, Akaike Information Criterion (AIC) values for logistic regression models trained using different sets of parameters. For each experiment, the AIC values for the 7 training settings have been ranked. The y-axis is the average ranking of each model over all GM12878 and K562 long-read TSS peaks from Cerberus (left) and LAPA (right), where logit[overlap TPM + DHS + peak_length] is the best model (i.e with the lowest AIC). e, Same-cell type ROC curves for logit[overlap TPM + DHS + peak_length]. Models tested on chr2 chr3 and trained on other chromosomes in the same cell line. f, Cross-cell type logit[overlap TPM + DHS + peak_length]. Distribution of AUROC values for long-read TSS experiments. Ex: To predict if a long-read TSS peak in K562 overlaps with a region in K562 RAMPAGE and CAGE in a cross-cell type manner, a model is trained on a GM12878.

**Figure S7.**
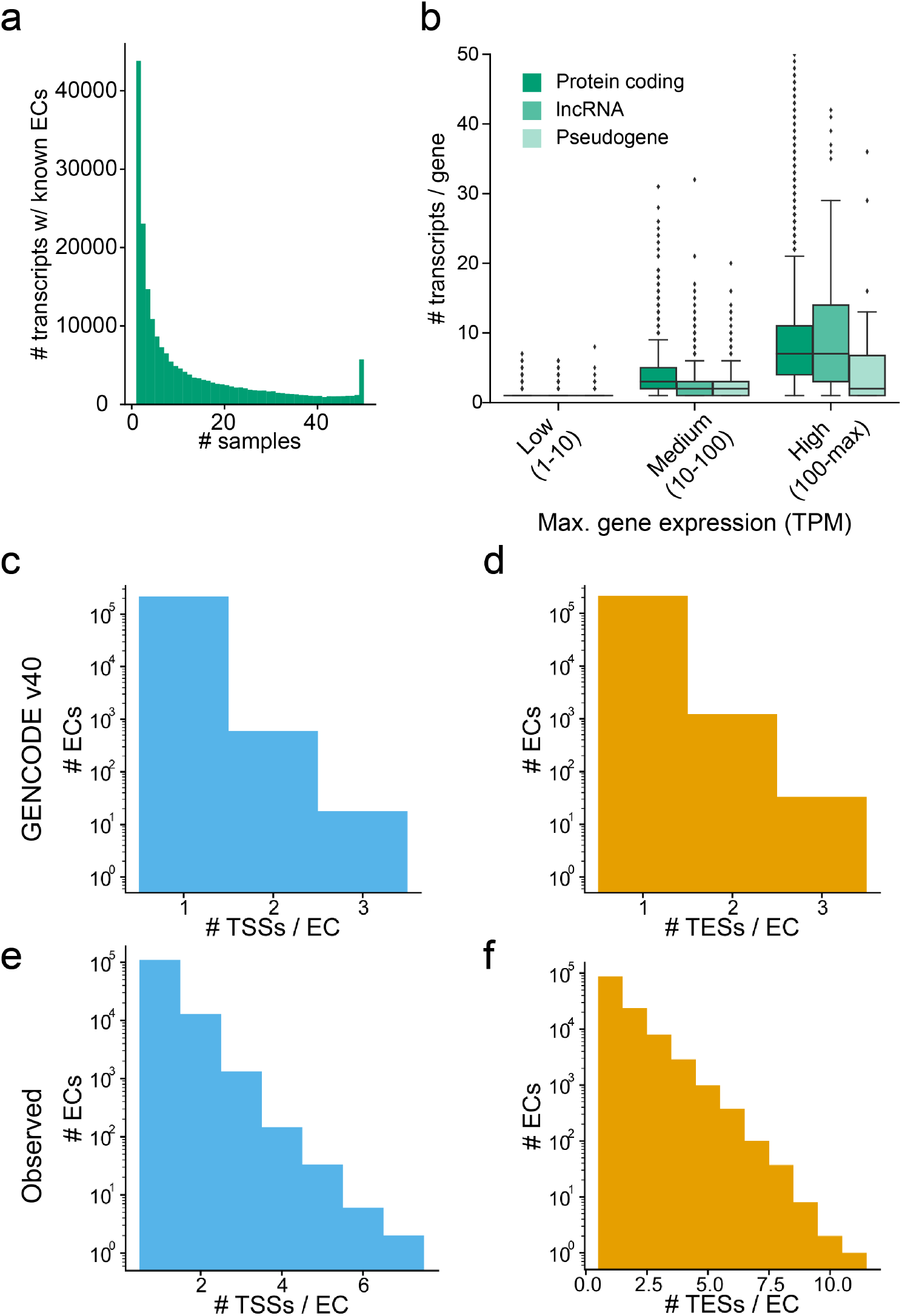
Characterization of observed transcripts from human LR-RNA-seq. a, Number of samples in which each transcript with a known EC is detected >= 1 TPM in the human ENCODE LR-RNA-seq dataset. b, Boxplot of, for the sample where a gene is most highly expressed, the number of transcripts expressed in that sample versus the TPM of the gene in that sample; split by gene biotype and gene expression bin. c-f, Number of unique TSSs or TESs per EC with at least 2 exons from transcripts c-d, annotated to polyA genes in GENCODE v40, e-f, detected >= 1 TPM from polyA genes in human ENCODE LR-RNA-seq.

**Figure S8.**
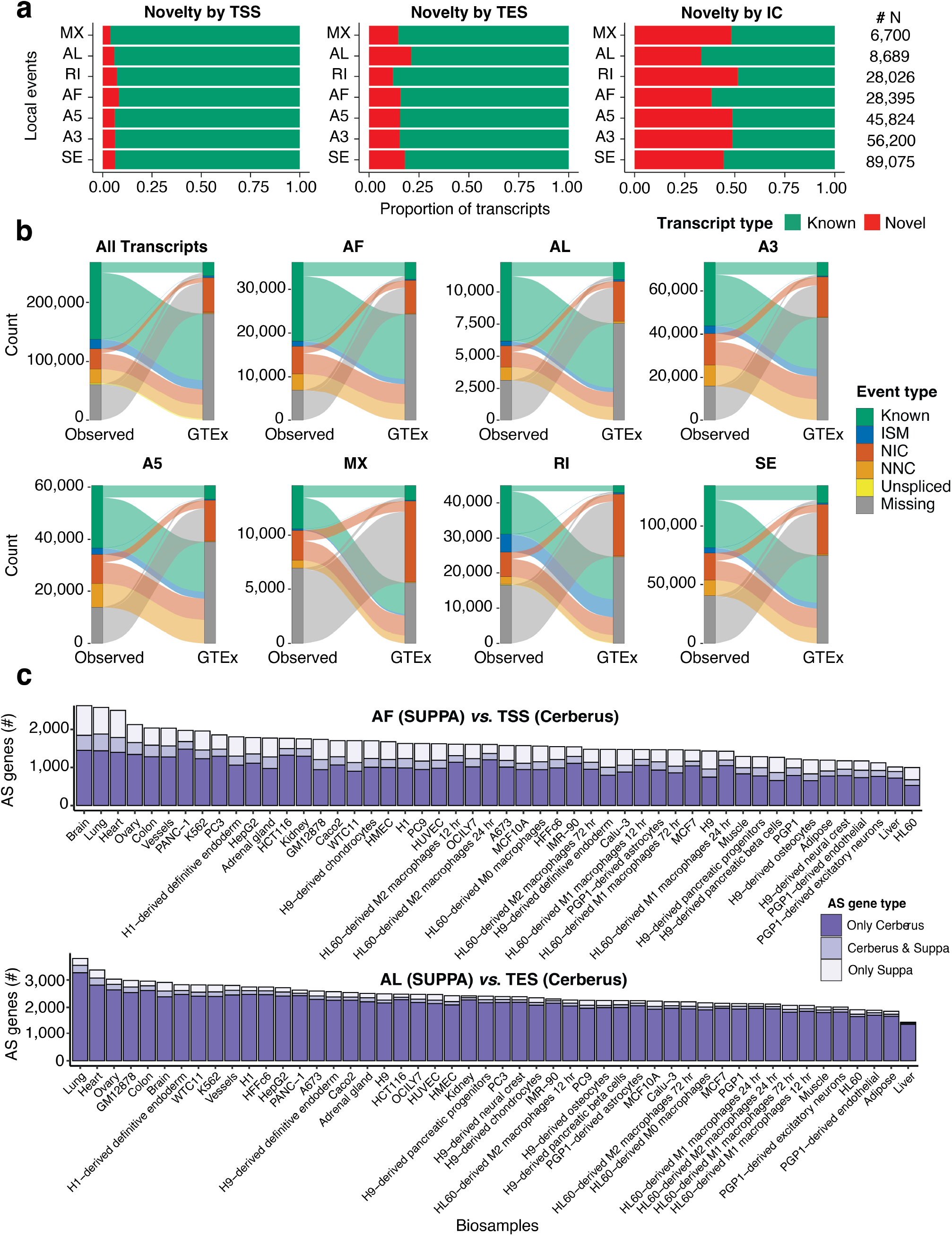
Alternative splicing (AS) detection by SUPPA and Cerberus. a, Barplots showing, for each type of local AS event detected by SUPPA (y axis), the proportion of observed known and novel transcripts identified by Cerberus (x axis), based on novel events at the TSS, TES, or EC. b, Sankey plots showing the number of transcripts classified as Known, Novel In Catalog (NIC), Novel Not in Catalog (NNC), Unspliced, or Missing by Cerberus and GTEx. In the first panel we show the numbers for all transcripts, while in the rest of the panels we focus on transcripts undergoing specific types of AS events. c, Barplot showing, for each LR-RNA-seq sample (x axis), the number of observed AS genes identified by both Cerberus and SUPPA (pink), only by Cerberus (dark gray), or only by SUPPA (light gray). In the upper panel we compare Alternative-First (AF)-AS genes by SUPPA and TSS-AS genes by Cerberus. In the lower panel we compare Alternative-Last (AL)-AS genes by SUPPA and TES-AS genes by Cerberus.

**Figure S9.**
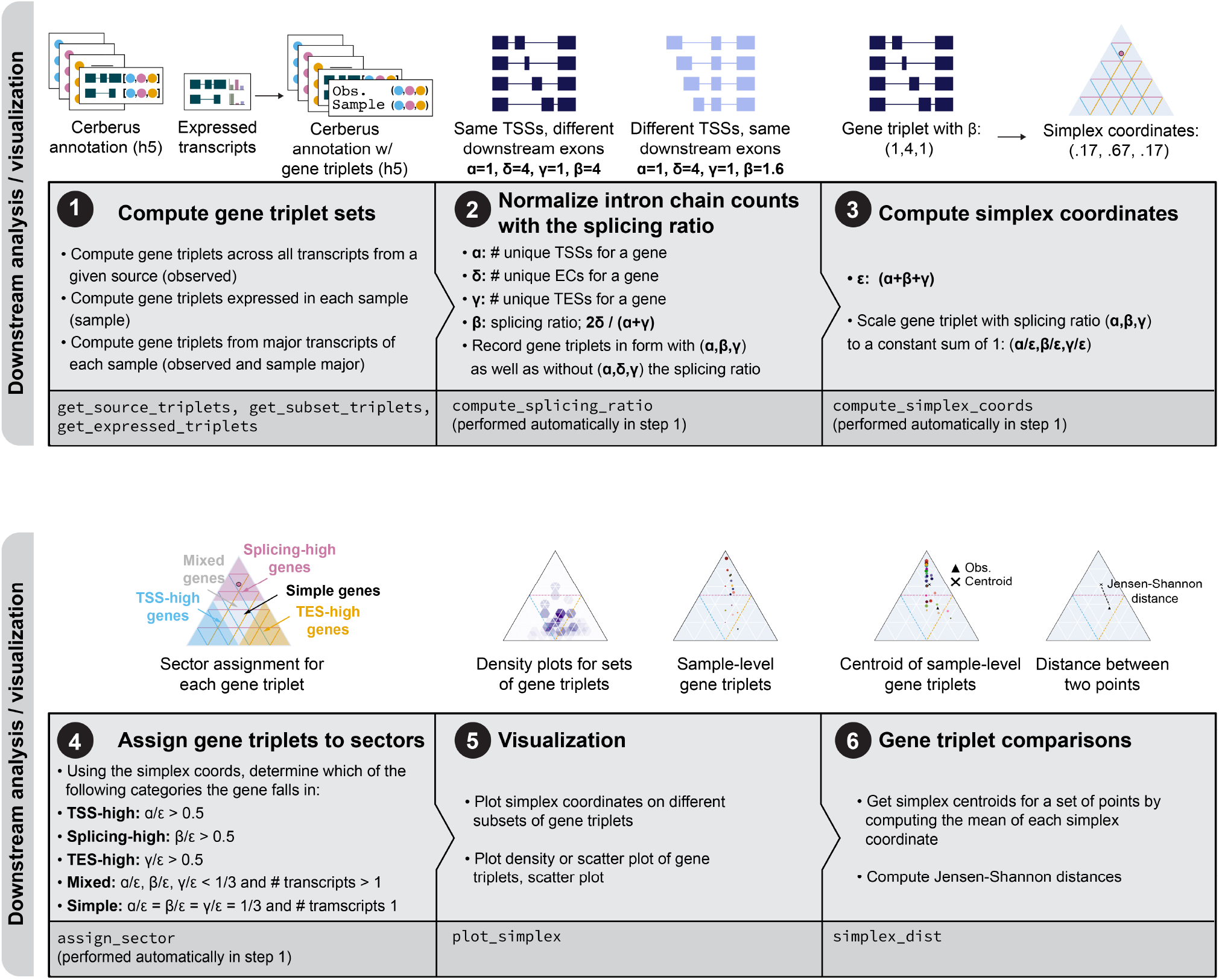
Overview of gene triplet based downstream analysis and visualization with Cerberus.

**Figure S10.**
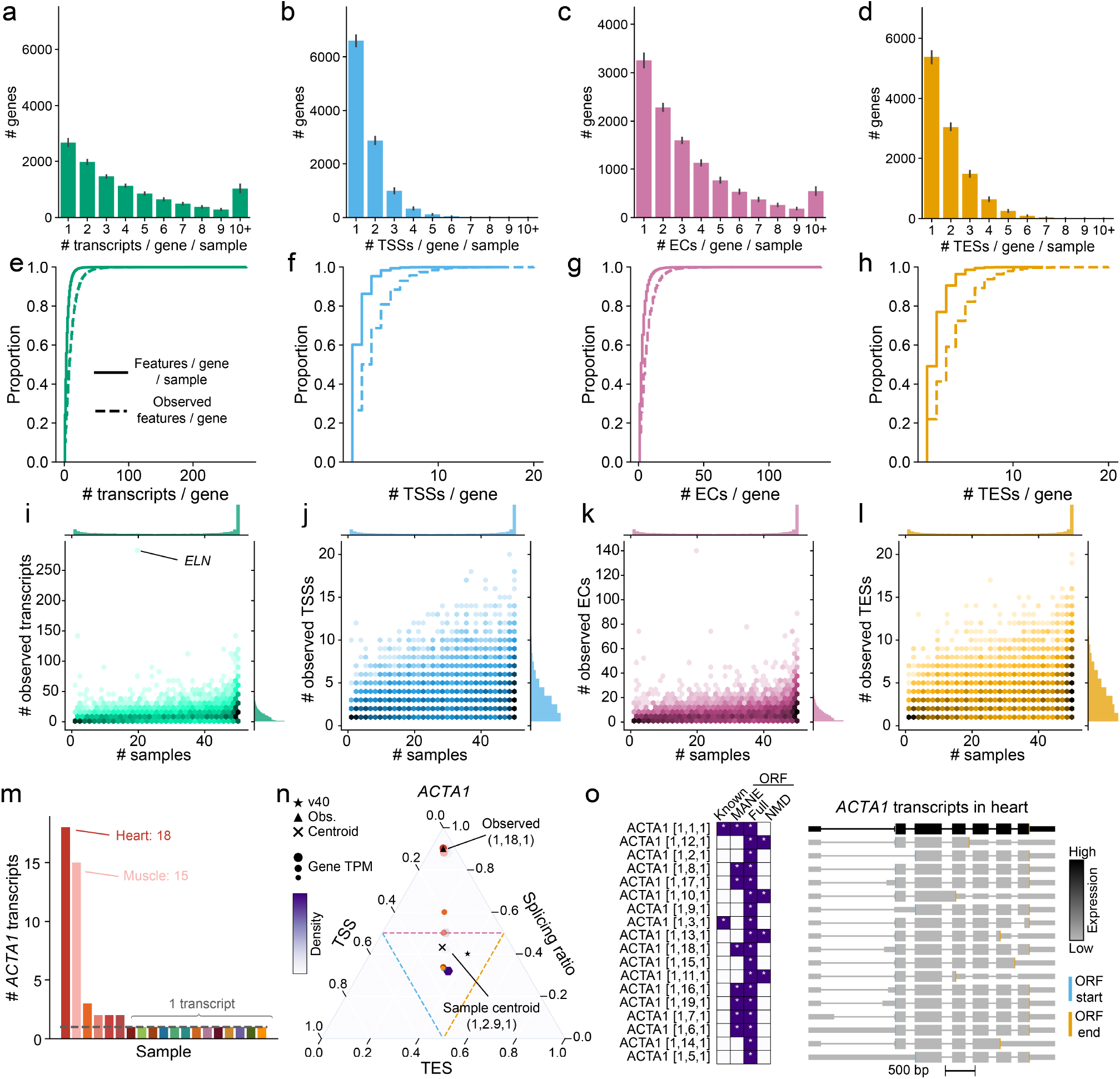
Uncovering sample-specific behavior of triplet features by comparing observed and sample-level gene triplets. a-d, Number of triplet features or transcripts detected per gene per sample for a, transcripts b, TSSs c, ECs d, TESs. e-h, Number of triplet features or transcripts per gene per sample and observed overall showing the proportion of the distribution that comes from each number of e, transcripts f, TSSs g, ECs h, TESs. i-l, Number of observed overall triplet features or transcripts per gene versus the number of samples each gene is expressed in for i, transcripts j, TSSs k, ECs l, TESs. m, Number of ACTA1 transcripts expressed in each sample. n, Gene structure simplex for ACTA1. Gene triplets with splicing ratio for the overall observed and sample-level centroid labeled. Simplex coordinates for the GENCODE v40, observed set, and centroid of the samples also shown for ACTA1. o, Browser models of transcripts of ACTA1 expressed >= 1 TPM in heart colored by expression level in TPM.

**Figure S11.**
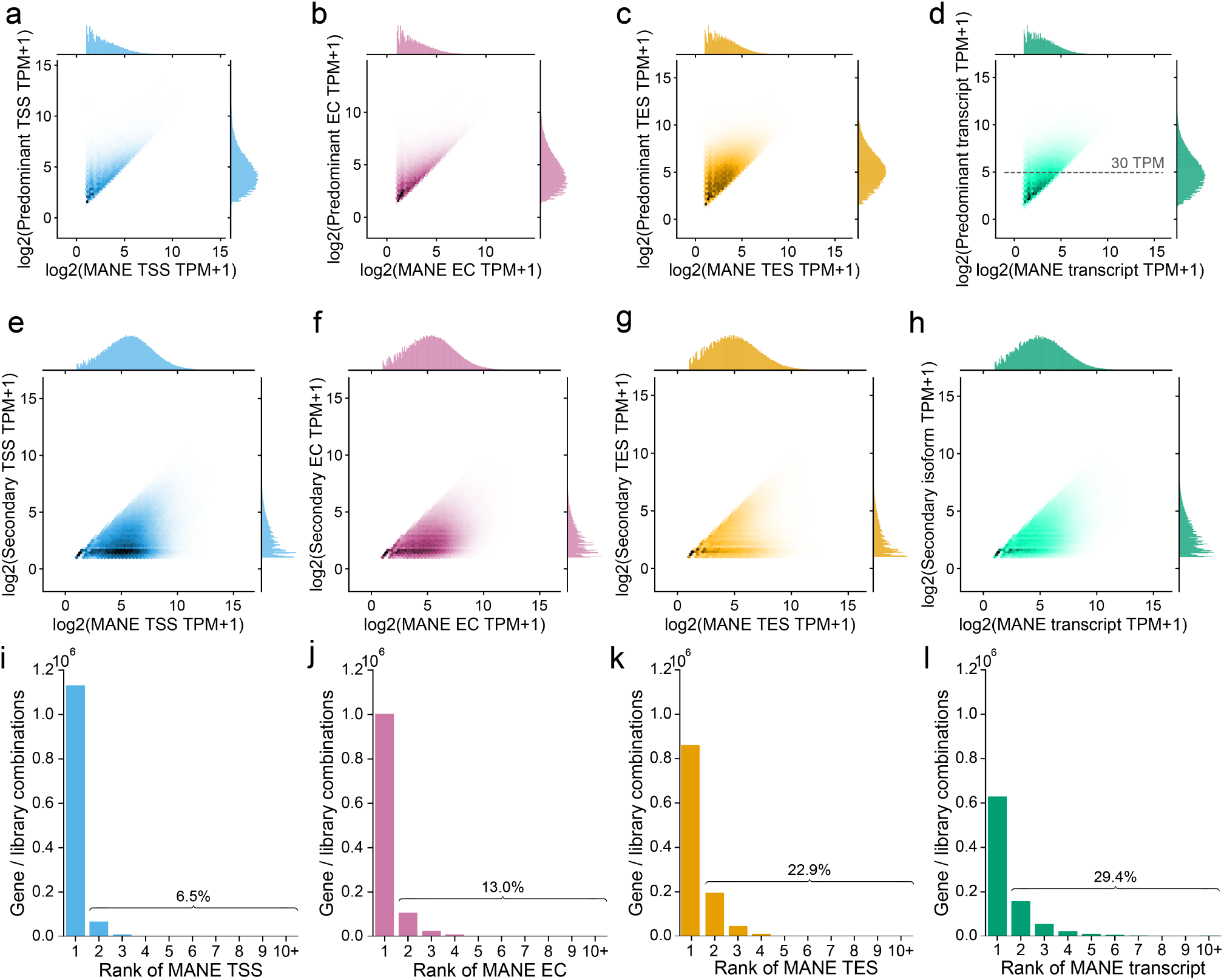
Rank and expression of predominant and MANE transcripts. a-d, For protein coding genes with MANE transcripts from GENCODE v40 where the predominant transcript or triplet feature is not the MANE transcript or triplet feature but is still expressed, expression of the predominant vs. the expression of the MANE a, TSS b, EC c, TES d, transcript. e-h, For genes where the MANE triplet feature or transcript is the predominant one and a secondary triplet feature or transcript is expressed, expression of the secondary vs. MANE e, TSS f, EC g, TES h, transcript. i-l, Rank of MANE i TSS j, EC k, TES l, transcript in each library where it is expressed.

**Figure S12.**
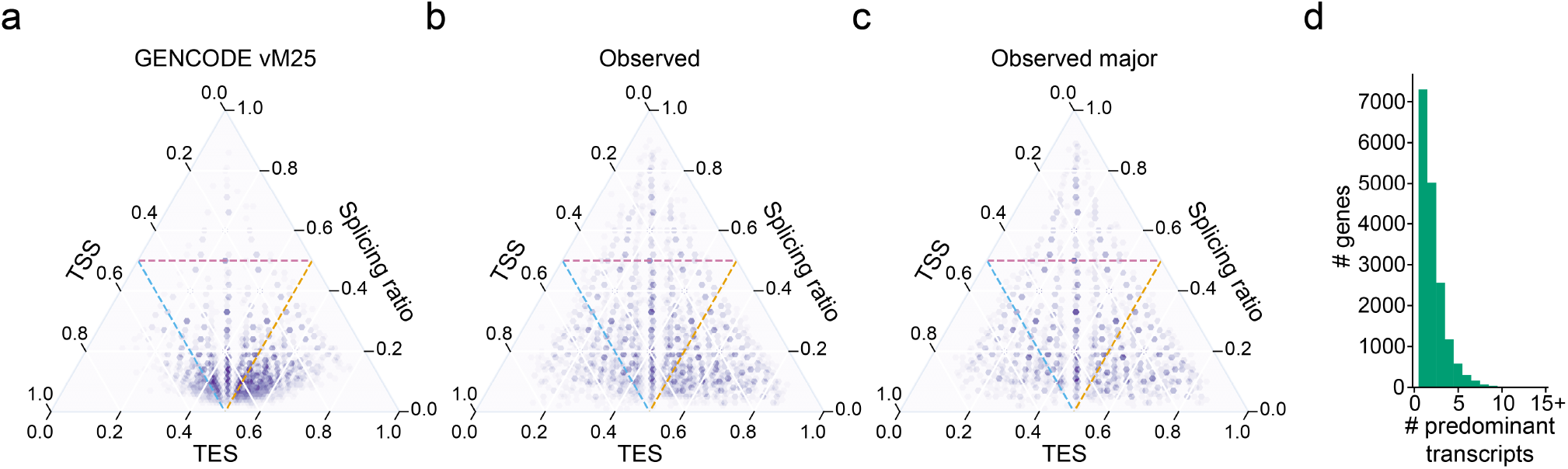
Rank and expression of predominant and MANE transcripts. a-c, Gene structure simplices for the transcripts from protein coding genes that are a, annotated in GENCODE vM25 where the parent gene is also detected in our mouse LR-RNA-seq dataset, b, the observed set of transcripts, those detected >= 1 TPM in the mouse ENCODE LR-RNA-seq dataset, c, the observed major set of transcripts, the union of major transcripts from each sample detected >= 1 TPM in the mouse ENCODE LR-RNA-seq dataset. d, Number of unique predominant transcripts detected >= 1 TPM across samples per gene.

## Notes

### Competing Interest Statement

The authors have declared no competing interest.

## Bibliography

1. Eddie Park, Zhicheng Pan, Zijun Zhang, Lan Lin, and Yi Xing. The Expanding Landscape of Alternative Splicing Variation in Human Populations. The American Journal of Human Genetics, 102(1):11–26, 2018. ISSN 0002-9297. doi: 10.1016/j.ajhg.2017.11.002.

2. Dafne Campigli Di Giammartino, Kensei Nishida, and James L. Manley. Mechanisms and Consequences of Alternative Polyadenylation. Molecular Cell, 43(6):853–866, 2011. ISSN 1097-2765. doi: 10.1016/j.molcel.2011.08.017.

3. Takeshi Ara, Fabrice Lopez, William Ritchie, Philippe Benech, and Daniel Gautheret. Conservation of alternative polyadenylation patterns in mammalian genes. BMC Genomics, 7 (1):189, 2006. doi: 10.1186/1471-2164-7-189.

4. Yi Xing and Christopher Lee. Alternative splicing and RNA selection pressure — evolutionary consequences for eukaryotic genomes. Nature Reviews Genetics, 7(7):499–509, 2006. ISSN 1471-0056. doi: 10.1038/nrg1896.

5. Hideki Nagasaki, Masanori Arita, Tatsuya Nishizawa, Makiko Suwa, and Osamu Gotoh. Species-specific variation of alternative splicing and transcriptional initiation in six eukaryotes. Gene, 364:53–62, 2005. ISSN 0378-1119. doi: 10.1016/j.gene.2005.07.027.

6. Nuala A. O’Leary, Mathew W. Wright, J. Rodney Brister, Stacy Ciufo, Diana Haddad, Rich McVeigh, Bhanu Rajput, Barbara Robbertse, Brian Smith-White, Danso Ako-Adjei, Alexander Astashyn, Azat Badretdin, Yiming Bao, Olga Blinkova, Vyacheslav Brover, Vyacheslav Chetvernin, Jinna Choi, Eric Cox, Olga Ermolaeva, Catherine M. Farrell, Tamara Goldfarb, Tripti Gupta, Daniel Haft, Eneida Hatcher, Wratko Hlavina, Vinita S. Joardar, Vamsi K. Kodali, Wenjun Li, Donna Maglott, Patrick Masterson, Kelly M. McGarvey, Michael R. Murphy, Kathleen O’Neill, Shashikant Pujar, Sanjida H. Rangwala, Daniel Rausch, Lillian D. Riddick, Conrad Schoch, Andrei Shkeda, Susan S. Storz, Hanzhen Sun, Francoise Thibaud-Nissen, Igor Tolstoy, Raymond E. Tully, Anjana R. Vatsan, Craig Wallin, David Webb, Wendy Wu, Melissa J. Landrum, Avi Kimchi, Tatiana Tatusova, Michael DiCuccio, Paul Kitts, Terence D. Murphy, and Kim D. Pruitt. Reference sequence (RefSeq) database at NCBI: current status, taxonomic expansion, and functional annotation. Nucleic Acids Research, 44(D1):D733–D745, 2016. ISSN 0305-1048. doi: 10.1093/nar/gkv1189.

7. Adam Frankish, Mark Diekhans, Irwin Jungreis, Julien Lagarde, Jane E Loveland, Jonathan M Mudge, Cristina Sisu, James C Wright, Joel Armstrong, If Barnes, Andrew Berry, Alexandra Bignell, Carles Boix, Silvia Carbonell Sala, Fiona Cunningham, Tomás Di Domenico, Sarah Donaldson, Ian T Fiddes, Carlos García Girón, Jose Manuel Gonzalez, Tiago Grego, Matthew Hardy, Thibaut Hourlier, Kevin L Howe, Toby Hunt, Osagie G Izuogu, Rory Johnson, Fergal J Martin, Laura Martínez, Shamika Mohanan, Paul Muir, Fabio C P Navarro, Anne Parker, Baikang Pei, Fernando Pozo, Ferriol Calvet Riera, Magali Ruffier, Bianca M Schmitt, Eloise Stapleton, Marie-Marthe Suner, Irina Sycheva, Barbara Uszczynska-Ratajczak, Maxim Y Wolf, Jinuri Xu, Yucheng T Yang, Andrew Yates, Daniel Zerbino, Yan Zhang, Jyoti S Choudhary, Mark Gerstein, Roderic Guigó, Tim J P Hubbard, Manolis Kellis, Benedict Paten, Michael L Tress, and Paul Flicek. GENCODE 2021. Nucleic Acids Research, 49(D1):gkaa1087–, 2020. ISSN 0305-1048. doi: 10.1093/nar/gkaa1087.

8. Joannella Morales, Shashikant Pujar, Jane E. Loveland, Alex Astashyn, Ruth Bennett, Andrew Berry, Eric Cox, Claire Davidson, Olga Ermolaeva, Catherine M. Farrell, Reham Fatima, Laurent Gil, Tamara Goldfarb, Jose M. Gonzalez, Diana Haddad, Matthew Hardy, Toby Hunt, John Jackson, Vinita S. Joardar, Michael Kay, Vamsi K. Kodali, Kelly M. McGarvey, Aoife McMahon, Jonathan M. Mudge, Daniel N. Murphy, Michael R. Murphy, Bhanu Rajput, Sanjida H. Rangwala, Lillian D. Riddick, Françoise Thibaud-Nissen, Glen Threadgold, Anjana R. Vatsan, Craig Wallin, David Webb, Paul Flicek, Ewan Birney, Kim D. Pruitt, Adam Frankish, Fiona Cunningham, and Terence D. Murphy. A joint NCBI and EMBL-EBI transcript set for clinical genomics and research. Nature, 604(7905):310–315, 2022. ISSN 0028-0836. doi: 10.1038/s41586-022-04558-8.

9. Barmak Modrek and Christopher Lee. A genomic view of alternative splicing. Nature Genetics, 30(1):13–19, 2002. ISSN 1061-4036. doi: 10.1038/ng0102-13.

10. Ali Mortazavi, Brian A Williams, Kenneth McCue, Lorian Schaeffer, and Barbara Wold. Mapping and quantifying mammalian transcriptomes by RNA-Seq. Nature Methods, 5(7):621–628, 2008. ISSN 1548-7091. doi: 10.1038/nmeth.1226.

11. Anthony Rhoads and Kin Fai Au. PacBio Sequencing and Its Applications. Genomics, Proteomics & Bioinformatics, 13(5):278–289, 2015. ISSN 1672-0229. doi: 10.1016/j.gpb.2015.08.002.

12. Daniel R Garalde, Elizabeth A Snell, Daniel Jachimowicz, Botond Sipos, Joseph H Lloyd, Mark Bruce, Nadia Pantic, Tigist Admassu, Phillip James, Anthony Warland, Michael Jordan, Jonah Ciccone, Sabrina Serra, Jemma Keenan, Samuel Martin, Luke McNeill E Jayne Wallace, Lakmal Jayasinghe, Chris Wright, Javier Blasco, Stephen Young, Denise Brocklebank, Sissel Juul, James Clarke, Andrew J Heron, and Daniel J Turner. Highly parallel direct RNA sequencing on an array of nanopores. Nature Methods, 15(3):201–206, 2018. ISSN 1548-7091. doi: 10.1038/nmeth.4577.

13. Ian Dunham, Anshul Kundaje, Shelley F. Aldred, Patrick J. Collins, Carrie A. Davis, Francis Doyle, Charles B. Epstein, Seth Frietze, Jennifer Harrow, Rajinder Kaul, Jainab Khatun, Bryan R. Lajoie, Stephen G. Landt, Bum-Kyu Lee, Florencia Pauli, Kate R. Rosenbloom, Peter Sabo, Alexias Safi, Amartya Sanyal, Noam Shoresh, Jeremy M. Simon, Lingyun Song, Nathan D. Trinklein, Robert C. Altshuler, Ewan Birney, James B. Brown, Chao Cheng, Sarah Djebali, Xianjun Dong, Ian Dunham, Jason Ernst, Terrence S. Furey, Mark Gerstein, Belinda Giardine, Melissa Greven, Ross C. Hardison, Robert S. Harris, Javier Herrero, Michael M. Hoffman, Sowmya Iyer, Manolis Kellis, Jainab Khatun, Pouya Kheradpour, Anshul Kundaje, Timo Lassmann, Qunhua Li, Xinying Lin, Georgi K. Marinov, Angelika Merkel, Ali Mortazavi, Stephen C. J. Parker, Timothy E. Reddy, Joel Rozowsky, Felix Schlesinger, Robert E. Thurman, Jie Wang, Lucas D. Ward, Troy W. Whitfield, Steven P. Wilder, Weisheng Wu, Hualin S. Xi, Kevin Y. Yip, Jiali Zhuang, Bradley E. Bernstein, Ewan Birney, Ian Dunham, Eric D. Green, Chris Gunter, Michael Snyder, Michael J. Pazin, Rebecca F. Lowdon, Laura A. L. Dillon, Leslie B. Adams, Caroline J. Kelly, Julia Zhang, Judith R. Wexler, Eric D. Green, Peter J. Good, Elise A. Feingold, Bradley E. Bernstein, Ewan Birney, Gregory E. Crawford, Job Dekker, Laura Elnitski, Peggy J. Farnham, Mark Gerstein, Morgan C. Giddings, Thomas R. Gingeras, Eric D. Green, Roderic Guigó, Ross C. Hardison, Timothy J. Hubbard, Manolis Kellis, W. James Kent, Jason D. Lieb, Elliott H. Margulies, Richard M. Myers, Michael Snyder, John A. Stamatoyannopoulos, Scott A. Tenenbaum, Zhiping Weng, Kevin P. White, Barbara Wold, Jainab Khatun, Yanbao Yu, John Wrobel, Brian A. Risk, Harsha P. Gunawardena, Heather C. Kuiper, Christopher W. Maier, Ling Xie, Xian Chen, Morgan C. Giddings, Bradley E. Bernstein, Charles B. Epstein, Noam Shoresh, Jason Ernst, Pouya Kheradpour, Tarjei S. Mikkelsen, Shawn Gillespie, Alon Goren, Oren Ram, Xiaolan Zhang, Li Wang, Robbyn Issner, Michael J. Coyne, Timothy Durham, Manching Ku, Thanh Truong, Lucas D. Ward, Robert C. Altshuler, Matthew L. Eaton, Manolis Kellis, Sarah Djebali, Carrie A. Davis, Angelika Merkel, Alex Dobin, Timo Lassmann, Ali Mortazavi, Andrea Tanzer, Julien Lagarde, Wei Lin, Felix Schlesinger, Chenghai Xue, Georgi K. Marinov, Jainab Khatun, Brian A. Williams, Chris Zaleski, Joel Rozowsky, Maik Röder, Felix Kokocinski, Rehab F. Abdelhamid, Tyler Alioto, Igor Antoshechkin, Michael T. Baer, Philippe Batut, Ian Bell, Kimberly Bell, Sudipto Chakrabortty, Xian Chen, Jacqueline Chrast, Joao Curado, Thomas Derrien, Jorg Drenkow, Erica Dumais, Jackie Dumais, Radha Duttagupta, Megan Fastuca, Kata Fejes-Toth, Pedro Ferreira, Sylvain Foissac, Melissa J. Fullwood, Hui Gao, David Gonzalez, Assaf Gordon, Harsha P. Gunawardena, Cédric Howald, Sonali Jha, Rory Johnson, Philipp Kapranov, Brandon King, Colin Kingswood, Guoliang Li, Oscar J. Luo, Eddie Park, Jonathan B. Preall, Kimberly Presaud, Paolo Ribeca, Brian A. Risk, Daniel Robyr, Xiaoan Ruan, Michael Sammeth, Kuljeet Singh Sandhu, Lorain Schaeffer, Lei-Hoon See, Atif Shahab, Jorgen Skancke, Ana Maria Suzuki, Hazuki Takahashi, Hagen Tilgner, Diane Trout, Nathalie Walters, Huaien Wang, John Wrobel, Yanbao Yu, Yoshihide Hayashizaki, Jennifer Harrow, Mark Gerstein, Timothy J. Hubbard, Alexandre Reymond, Stylianos E. Antonarakis, Gregory J. Hannon, Morgan C. Giddings, Yijun Ruan, Barbara Wold, Piero Carninci, Roderic Guigó, Thomas R. Gingeras, Kate R. Rosenbloom, Cricket A. Sloan, Katrina Learned, Venkat S. Malladi, Matthew C. Wong, Galt P. Barber, Melissa S. Cline, Timothy R. Dreszer, Steven G. Heitner, Donna Karolchik, W. James Kent, Vanessa M. Kirkup, Laurence R. Meyer, Jeffrey C. Long, Morgan Maddren, Brian J. Raney, Terrence S. Furey, Lingyun Song, Linda L. Grasfeder, Paul G. Giresi, Bum-Kyu Lee, Anna Battenhouse, Nathan C. Sheffield, Jeremy M. Simon, Kimberly A. Showers, Alexias Safi, Darin London, Akshay A. Bhinge, Christopher Shestak, Matthew R. Schaner, Seul Ki Kim, Zhuzhu Z. Zhang, Piotr A. Mieczkowski, Joanna O. Mieczkowska, Zheng Liu, Ryan M. McDaniell, Yunyun Ni, Naim U. Rashid, Min Jae Kim, Sheera Adar, Zhancheng Zhang, Tianyuan Wang, Deborah Winter, Damian Keefe, Ewan Birney, Vishwanath R. Iyer, Jason D. Lieb, Gregory E. Crawford, Guoliang Li, Kuljeet Singh Sandhu, Meizhen Zheng, Ping Wang, Oscar J. Luo, Atif Shahab, Melissa J. Fullwood, Xiaoan Ruan, Yijun Ruan, Richard M. Myers, Florencia Pauli, Brian A. Williams, Jason Gertz, Georgi K. Marinov, Timothy E. Reddy, Jost Vielmetter, E. Partridge, Diane Trout, Katherine E. Varley, Clarke Gasper, Anita Bansal, Shirley Pepke, Preti Jain, Henry Amrhein, Kevin M. Bowling, Michael Anaya, Marie K. Cross, Brandon King, Michael A. Muratet, Igor Antoshechkin, Kimberly M. Newberry, Kenneth McCue, Amy S. Nesmith, Katherine I. Fisher-Aylor, Barbara Pusey, Gilberto DeSalvo, Stephanie L. Parker, Sreeram Balasubramanian, Nicholas S. Davis, Sarah K. Meadows, Tracy Eggleston, Chris Gunter, J. Scott Newberry, Shawn E. Levy, Devin M. Absher, Ali Mortazavi, Wing H. Wong, Barbara Wold, Matthew J. Blow, Axel Visel, Len A. Pennachio, Laura Elnitski, Elliott H. Margulies, Stephen C. J. Parker, Hanna M. Petrykowska, Alexej Abyzov, Bronwen Aken, Daniel Barrell, Gemma Barson, Andrew Berry, Alexandra Bignell, Veronika Boychenko, Giovanni Bussotti, Jacqueline Chrast, Claire Davidson, Thomas Derrien, Gloria Despacio-Reyes, Mark Diekhans, Iakes Ezkurdia, Adam Frankish, James Gilbert, Jose Manuel Gonzalez, Ed Griffiths, Rachel Harte, David A. Hendrix, Cédric Howald, Toby Hunt, Irwin Jungreis, Mike Kay, Ekta Khurana, Felix Kokocinski, Jing Leng, Michael F. Lin, Jane Loveland, Zhi Lu, Deepa Manthravadi, Marco Mariotti, Jonathan Mudge, Gaurab Mukherjee, Cedric Notredame, Baikang Pei, Jose Manuel Rodriguez, Gary Saunders, Andrea Sboner, Stephen Searle, Cristina Sisu, Catherine Snow, Charlie Steward, Andrea Tanzer, Electra Tapanari, Michael L. Tress, Marijke J. van Baren, Nathalie Walters, Stefan Washietl, Laurens Wilming, Amonida Zadissa, Zhengdong Zhang, Michael Brent, David Haussler, Manolis Kellis, Alfonso Valencia, Mark Gerstein, Alexandre Reymond, Roderic Guigó, Jennifer Harrow, Timothy J. Hubbard, Stephen G. Landt, Seth Frietze, Alexej Abyzov, Nick Addleman, Roger P. Alexander, Raymond K. Auerbach, Suganthi Balasubramanian, Keith Bettinger, Nitin Bhardwaj, Alan P. Boyle, Alina R. Cao, Philip Cayting, Alexandra Charos, Yong Cheng, Chao Cheng, Catharine Eastman, Ghia Euskirchen, Joseph D. Fleming, Fabian Grubert, Lukas Habegger, Manoj Hariharan, Arif Harmanci, Sushma Iyengar, Victor X. Jin, Konrad J. Karczewski, Maya Kasowski, Phil Lacroute, Hugo Lam, Nathan Lamarre-Vincent, Jing Leng, Jin Lian, Marianne Lindahl-Allen, Renqiang Min, Benoit Miotto, Hannah Monahan, Zarmik Mo-qtaderi, Xinmeng J. Mu, Henriette O’Geen, Zhengqing Ouyang, Dorrelyn Patacsil, Baikang Pei, Debasish Raha, Lucia Ramirez, Brian Reed, Joel Rozowsky, Andrea Sboner, Minyi Shi, Cristina Sisu, Teri Slifer, Heather Witt, Linfeng Wu, Xiaoqin Xu, Koon-Kiu Yan, Xinqiong Yang, Kevin Y. Yip, Zhengdong Zhang, Kevin Struhl, Sherman M. Weissman, Mark Gerstein, Peggy J. Farnham, Michael Snyder, Scott A. Tenenbaum, Luiz O. Penalva, Francis Doyle, Subhradip Karmakar, Stephen G. Landt, Raj R. Bhanvadia, Alina Choudhury, Marc Domanus, Lijia Ma, Jennifer Moran, Dorrelyn Patacsil, Teri Slifer, Alec Victorsen, Xinqiong Yang, Michael Snyder, Kevin P. White, Thomas Auer, Lazaro Centanin, Michael Eichenlaub, Franziska Gruhl, Stephan Heermann, Burkhard Hoeckendorf, Daigo Inoue, Tanja Kellner, Stephan Kirchmaier, Claudia Mueller, Robert Reinhardt, Lea Schertel, Stephanie Schneider, Rebecca Sinn, Beate Wittbrodt, Jochen Wittbrodt, Zhiping Weng, Troy W. Whitfield, Jie Wang, Patrick J. Collins, Shelley F. Aldred, Nathan D. Trinklein, E. Christopher Partridge, Richard M. Myers, Job Dekker, Gaurav Jain, Bryan R. Lajoie, Amartya Sanyal, Gayathri Balasundaram, Daniel L. Bates, Rachel Byron, Theresa K. Canfield, Morgan J. Diegel, Douglas Dunn, Abigail K. Ebersol, Tristan Frum, Kavita Garg, Erica Gist, R. Scott Hansen, Lisa Boatman, Eric Haugen, Richard Humbert, Gaurav Jain, Audra K. Johnson, Ericka M. Johnson, Tattyana V. Kutyavin, Bryan R. Lajoie, Kristen Lee, Dimitra Lotakis, Matthew T. Maurano, Shane J. Neph, Fiedencio V. Neri, Eric D. Nguyen, Hongzhu Qu, Alex P. Reynolds, Vaughn Roach, Eric Rynes, Peter Sabo, Minerva E. Sanchez, Richard S. Sandstrom, Amartya Sanyal, Anthony O. Shafer, Andrew B. Stergachis, Sean Thomas, Robert E. Thurman, Benjamin Vernot, Jeff Vierstra, Shinny Vong, Hao Wang, Molly A. Weaver, Yongqi Yan, Miaohua Zhang, Joshua M. Akey, Michael Bender, Michael O. Dorschner, Mark Groudine, Michael J. MacCoss, Patrick Navas, George Stamatoyannopoulos, Rajinder Kaul, Job Dekker, John A. Stamatoyannopoulos, Ian Dunham, Kathryn Beal, Alvis Brazma, Paul Flicek, Javier Herrero, Nathan Johnson, Damian Keefe, Margus Lukk, Nicholas M. Luscombe, Daniel Sobral, Juan M. Vaquerizas, Steven P. Wilder, Serafim Batzoglou, Arend Sidow, Nadine Hussami, Sofia Kyriazopoulou-Panagiotopoulou, Max W. Libbrecht, Marc A. Schaub, Anshul Kundaje, Ross C. Hardison, Webb Miller, Belinda Giardine, Robert S. Harris, Weisheng Wu, Peter J. Bickel, Balazs Banfai, Nathan P. Boley, James B. Brown, Haiyan Huang, Qunhua Li, Jingyi Jessica Li, William Stafford Noble, Jeffrey A. Bilmes, Orion J. Buske, Michael M. Hoffman, Avinash D. Sahu, Peter V. Kharchenko, Peter J. Park, Dannon Baker, James Taylor, Zhiping Weng, Sowmya Iyer, Xianjun Dong, Melissa Greven, Xinying Lin, Jie Wang, Hualin S. Xi, Jiali Zhuang, Mark Gerstein, Roger P. Alexander, Suganthi Balasubramanian, Chao Cheng, Arif Harmanci, Lucas Lochovsky, Renqiang Min, Xinmeng J. Mu, Joel Rozowsky, Koon-Kiu Yan, Kevin Y. Yip, and Ewan Birney. An integrated encyclopedia of DNA elements in the human genome. Nature, 489(7414):57–74, 2012. ISSN 0028-0836. doi: 10.1038/nature11247.

14. Sarah Djebali, Carrie A. Davis, Angelika Merkel, Alex Dobin, Timo Lassmann, Ali Mortazavi, Andrea Tanzer, Julien Lagarde, Wei Lin, Felix Schlesinger, Chenghai Xue, Georgi K. Marinov, Jainab Khatun, Brian A. Williams, Chris Zaleski, Joel Rozowsky, Maik Röder, Felix Kokocinski, Rehab F. Abdelhamid, Tyler Alioto, Igor Antoshechkin, Michael T. Baer, Nadav S. Bar, Philippe Batut, Kimberly Bell, Ian Bell, Sudipto Chakrabortty, Xian Chen, Jacqueline Chrast, Joao Curado, Thomas Derrien, Jorg Drenkow, Erica Dumais, Jacqueline Dumais, Radha Duttagupta, Emilie Falconnet, Meagan Fastuca, Kata Fejes-Toth, Pedro Ferreira, Sylvain Foissac, Melissa J. Fullwood, Hui Gao, David Gonzalez, Assaf Gordon, Harsha Gunawardena, Cedric Howald, Sonali Jha, Rory Johnson, Philipp Kapranov, Brandon King, Colin Kingswood, Oscar J. Luo, Eddie Park, Kimberly Persaud, Jonathan B. Preall, Paolo Ribeca, Brian Risk, Daniel Robyr, Michael Sammeth, Lorian Schaffer, Lei-Hoon See, Atif Shahab, Jorgen Skancke, Ana Maria Suzuki, Hazuki Takahashi, Hagen Tilgner, Diane Trout, Nathalie Walters, Huaien Wang, John Wrobel, Yanbao Yu, Xiaoan Ruan, Yoshihide Hayashizaki, Jennifer Harrow, Mark Gerstein, Tim Hubbard, Alexandre Reymond, Stylianos E. Antonarakis, Gregory Hannon, Morgan C. Giddings, Yijun Ruan, Barbara Wold, Piero Carninci, Roderic Guigó, and Thomas R. Gingeras. Landscape of transcription in human cells. Nature, 489(7414):101–108, 2012. ISSN 0028-0836. doi: 10.1038/nature11233.

15. Federico Abascal, Reyes Acosta, Nicholas J. Addleman, Jessika Adrian, Veena Afzal, Rizi Ai, Bronwen Aken, Jennifer A. Akiyama, Omar Al Jammal, Henry Amrhein, Stacie M. Anderson, Gregory R. Andrews, Igor Antoshechkin, Kristin G. Ardlie, Joel Armstrong, Matthew Astley, Budhaditya Banerjee, Amira A. Barkal, If H. A. Barnes, Iros Barozzi, Daniel Barrell, Gemma Barson, Daniel Bates, Ulugbek K. Baymuradov, Cassandra Bazile, Michael A. Beer, Samantha Beik, M. A. Bender, Ruth Bennett, Louis Philip Benoit Bouvrette, Bradley E. Bernstein, Andrew Berry, Anand Bhaskar, Alexandra Bignell, Steven M. Blue, David M. Bodine, Carles Boix, Nathan Boley, Tyler Borrman, Beatrice Borsari, Alan P. Boyle, Laurel A. Brandsmeier, Alessandra Breschi, Emery H. Bresnick, Jason A. Brooks, Michael Buckley, Christopher B. Burge, Rachel Byron, Eileen Cahill, Lingling Cai, Lulu Cao, Mark Carty, Rosa G. Castanon, Andres Castillo, Hassan Chaib, Esther T. Chan, Daniel R. Chee, Sora Chee, Hao Chen, Huaming Chen, Jia-Yu Chen, Songjie Chen, J. Michael Cherry, Surya B. Chhetri, Jyoti S. Choudhary, Jacqueline Chrast, Dongjun Chung, Declan Clarke, Neal A. L. Cody, Candice J. Coppola, Julie Coursen Anthony M. D’Ippolito, Stephen Dalton, Cassidy Danyko, Claire Davidson, Jose Davila-Velderrain, Carrie A. Davis, Job Dekker, Alden Deran, Gilberto DeSalvo, Gloria Despacio-Reyes, Colin N. Dewey, Diane E. Dickel, Morgan Diegel, Mark Diekhans, Vishnu Dileep, Bo Ding, Sarah Djebali, Alexander Dobin, Daniel Dominguez, Sarah Donaldson, Jorg Drenkow, Timothy R. Dreszer, Yotam Drier, Michael O. Duff, Douglass Dunn, Catharine Eastman, Joseph R. Ecker, Matthew D. Edwards, Nicole El-Ali, Shaimae I. Elhajjajy, Keri Elkins, Andrew Emili, Charles B. Epstein, Rachel C. Evans, Iakes Ezkurdia, Kaili Fan, Peggy J. Farnham, Nina P. Farrell, Elise A. Feingold, Anne-Maud Ferreira, Katherine Fisher-Aylor, Stephen Fitzgerald, Paul Flicek, Chuan Sheng Foo, Kevin Fortier, Adam Frankish, Peter Freese, Shaliu Fu, Xiang-Dong Fu, Yu Fu, Yoko Fukuda-Yuzawa, Mariateresa Fulciniti, Alister P. W. Funnell, Idan Gabdank, Timur Galeev, Ming-shi Gao, Carlos Garcia Giron, Tyler H. Garvin, Chelsea Anne Gelboin-Burkhart, Grigorios Georgolopoulos, Mark B. Gerstein, Belinda M. Giardine, David K. Gifford, David M. Gilbert, Daniel A. Gilchrist, Shawn Gillespie, Thomas R. Gingeras, Peng Gong, Alvaro Gonzalez, Jose M. Gonzalez, Peter Good, Alon Goren, David U. Gorkin, Brenton R. Graveley, Michael Gray, Jack F. Greenblatt, Ed Griffiths, Mark T. Groudine, Fabian Grubert, Mengting Gu, Roderic Guigó, Hongbo Guo, Yu Guo, Yuchun Guo, Gamze Gursoy, Maria Gutierrez-Arcelus, Jessica Halow, Ross C. Hardison, Matthew Hardy, Manoj Hariharan, Arif Harmanci, Anne Harrington, Jennifer L. Harrow, Tatsunori B. Hashimoto, Richard D. Hasz, Meital Hatan, Eric Haugen, James E. Hayes, Peng He, Yupeng He, Nastaran Heidari, David Hendrickson, Elisabeth F. Heuston, Jason A. Hilton, Benjamin C. Hitz, Abigail Hochman, Cory Holgren, Lei Hou, Shuyu Hou, Yun-Hua E. Hsiao, Shanna Hsu, Hui Huang, Tim J. Hubbard, Jack Huey, Timothy R. Hughes, Toby Hunt, Sean Ibarrientos, Robbyn Issner, Mineo Iwata, Osagie Izuogu, Tommi Jaakkola, Nader Jameel, Camden Jansen, Lixia Jiang, Peng Jiang, Audra Johnson, Rory Johnson, Irwin Jungreis, Madhura Kadaba, Maya Kasowski, Mary Kasparian, Momoe Kato, Rajinder Kaul, Trupti Kawli, Michael Kay, Judith C. Keen, Sunduz Keles, Cheryl A. Keller, David Kelley, Manolis Kellis, Pouya Kheradpour, Daniel Sunwook Kim, Anthony Kirilusha, Robert J. Klein, Birgit Knoechel, Samantha Kuan, Michael J. Kulik, Sushant Kumar, Anshul Kundaje, Tanya Kutyavin, Julien Lagarde, Bryan R. Lajoie, Nicole J. Lambert, John Lazar, Ah Young Lee, Donghoon Lee, Elizabeth Lee, Jin Wook Lee, Kristen Lee, Christina S. Leslie, Shawn Levy, Bin Li, Hairi Li, Nan Li, Shantao Li, Xiangrui Li, Yang I. Li, Ying Li, Yining Li, Yue Li, Jin Lian, Maxwell W. Libbrecht, Shin Lin, Yiing Lin, Dianbo Liu, Jason Liu, Peng Liu, Tingting Liu, X. Shirley Liu, Yan Liu, Yaping Liu, Maria Long, Shaoke Lou, Jane Loveland, Aiping Lu, Yuheng Lu, Eric Lécuyer, Lijia Ma, Mark Mackiewicz, Brandon J. Mannion, Michael Mannstadt, Deepa Manthravadi, Georgi K. Marinov, Fergal J. Martin, Eugenio Mattei, Kenneth McCue, Megan McEown, Graham McVicker, Sarah K. Meadows, Alex Meissner, Eric M. Mendenhall, Christopher L. Messer, Wouter Meuleman, Clifford Meyer, Steve Miller, Matthew G. Milton, Tejaswini Mishra, Dianna E. Moore, Helen M. Moore, Jill E. Moore, Samuel H. Moore, Jennifer Moran, Ali Mortazavi, Jonathan M. Mudge, Nikhil Munshi, Rabi Murad, Richard M. Myers, Vivek Nandakumar, Preetha Nandi, Anil M. Narasimha, Aditi K. Narayanan, Hannah Naughton, Fabio C. P. Navarro, Patrick Navas, Jurijs Nazarovs, Jemma Nelson, Shane Neph, Fidencio Jun Neri, Joseph R. Nery, Amy R. Nesmith, J. Scott Newberry, Kimberly M. Newberry, Vu Ngo, Rosy Nguyen, Thai B. Nguyen, Tung Nguyen, Andrew Nishida, William S. Noble, Catherine S. Novak, Eva Maria Novoa, Briana Nuñez, Charles W. O’Donnell, Sara Olson, Kathrina C. Onate, Ericka Otterman, Hakan Ozadam, Michael Pagan, Tsultrim Palden, Xinghua Pan, Yongjin Park, E. Christopher Partridge, Benedict Paten, Florencia Pauli-Behn, Michael J. Pazin, Baikang Pei, Len A. Pennacchio, Alexander R. Perez, Emily H. Perry, Dmitri D. Pervouchine, Nishigandha N. Phalke, Quan Pham, Doug H. Phanstiel, Ingrid Plajzer-Frick, Gabriel A. Pratt, Henry E. Pratt, Sebastian Preissl, Jonathan K. Pritchard, Yuri Pritykin, Michael J. Purcaro, Qian Qin, Giovanni Quinones-Valdez, Ines Rabano, Ernest Radovani, Anil Raj, Nisha Rajagopal, Oren Ram, Lucia Ramirez, Ricardo N. Ramirez, Dylan Rausch, Soumya Raychaudhuri, Joseph Raymond, Rozita Razavi, Timothy E. Reddy, Thomas M. Reimonn, Bing Ren, Alexandre Reymond, Alex Reynolds, Suhn K. Rhie, John Rinn, Miguel Rivera, Juan Carlos Rivera-Mulia, Brian S. Roberts, Jose Manuel Rodriguez, Joel Rozowsky, Russell Ryan, Eric Rynes, Denis N. Salins, Richard Sandstrom, Takayo Sasaki, Shashank Sathe, Daniel Savic, Alexandra Scavelli, Jonathan Scheiman, Christoph Schlaffner, Jeffery A. Schloss, Frank W. Schmitges, Lei Hoon See, Anurag Sethi, Manu Setty, Anthony Shafer, Shuo Shan, Eilon Sharon, Quan Shen, Yin Shen, Richard I. Sherwood, Minyi Shi, Sunyoung Shin, Noam Shoresh, Kyle Siebenthall, Cristina Sisu, Teri Slifer, Cricket A. Sloan, Anna Smith, Valentina Snetkova, Michael P. Snyder, Damek V. Spacek, Sharanya Srinivasan, Rohith Srivas, George Stamatoyannopoulos, John A. Stamatoyannopoulos, Rebecca Stanton, Dave Steffan, Sandra Stehling-Sun, J. Seth Strattan, Amanda Su, Balaji Sundararaman, Marie-Marthe Suner, Tahin Syed, Matt Szynkarek, Forrest Y. Tanaka, Danielle Tenen, Mingxiang Teng, Jeffrey A. Thomas, Dave Toffey, Michael L. Tress, Diane E. Trout, Gosia Trynka, Junko Tsuji, Sean A. Upchurch, Oana Ursu, Barbara Uszczynska-Ratajczak, Mia C. Uziel, Alfonso Valencia, Benjamin Van Biber, Arjan G. van der Velde, Eric L. Van Nostrand, Yekaterina Vaydylevich, Jesus Vazquez, Alec Victorsen, Jost Vielmetter, Jeff Vierstra, Axel Visel, Anna Vlasova, Christopher M. Vockley, Simona Volpi, Shinny Vong, Hao Wang, Mengchi Wang, Qin Wang, Ruth Wang, Tao Wang, Wei Wang, Xiaofeng Wang, Yanli Wang, Nathaniel K. Watson, Xintao Wei, Zhijie Wei, Hendrik Weisser, Sherman M. Weissman, Rene Welch, Robert E. Welikson, Zhiping Weng, Harm-Jan Westra, John W. Whitaker, Collin White, Kevin P. White, Andre Wildberg, Brian A. Williams, David Wine, Heather N. Witt, Barbara Wold, Maxim Wolf, James Wright, Rui Xiao, Xinshu Xiao, Jie Xu, Jinrui Xu, Koon-Kiu Yan, Yongqi Yan, Hongbo Yang, Xinqiong Yang, Yi-Wen Yang, Galip Gürkan Yardimci, Brian A. Yee, Gene W. Yeo, Taylor Young, Tianxiong Yu, Feng Yue, Chris Zaleski, Chongzhi Zang, Haoyang Zeng, Weihua Zeng, Daniel R. Zerbino, Jie Zhai, Lijun Zhan, Ye Zhan, Bo Zhang, Jialing Zhang, Jing Zhang, Kai Zhang, Lijun Zhang, Peng Zhang, Qi Zhang, Xiao-Ou Zhang, Yanxiao Zhang, Zhizhuo Zhang, Yuan Zhao, Ye Zheng, Guoqing Zhong, Xiao-Qiao Zhou, Yun Zhu, Jared Zimmerman, Jill E. Moore, Michael J. Purcaro, Henry E. Pratt, Charles B. Epstein, Noam Shoresh, Jessika Adrian, Trupti Kawli, Carrie A. Davis, Alexander Dobin, Rajinder Kaul, Jessica Halow, Eric L. Van Nostrand, Peter Freese, David U. Gorkin, Yin Shen, Yupeng He, Mark Mackiewicz, Florencia Pauli-Behn, Brian A. Williams, Ali Mortazavi, Cheryl A. Keller, Xiao-Ou Zhang, Shaimae I. Elhajjajy, Jack Huey, Diane E. Dickel, Valentina Snetkova, Xintao Wei, Xiaofeng Wang, Juan Carlos Rivera-Mulia, Joel Rozowsky, Jing Zhang, Surya B. Chhetri, Jialing Zhang, Alec Victorsen, Kevin P. White, Axel Visel, Gene W. Yeo, Christopher B. Burge, Eric Lécuyer, David M. Gilbert, Job Dekker, John Rinn, Eric M. Mendenhall, Joseph R. Ecker, Manolis Kellis, Robert J. Klein, William S. Noble, Anshul Kundaje, Roderic Guigó, Peggy J. Farnham, J. Michael Cherry, Richard M. Myers, Bing Ren, Brenton R. Graveley, Mark B. Gerstein, Len A. Pennacchio, Michael P. Snyder, Bradley E. Bernstein, Barbara Wold, Ross C. Hardison, Thomas R. Gingeras, John A. Stamatoyannopoulos, and Zhiping Weng. Expanded encyclopaedias of DNA elements in the human and mouse genomes. Nature, 583(7818):699–710, 2020. ISSN 0028-0836. doi: 10.1038/s41586-020-2493-4.

16. Marina Lizio, Jayson Harshbarger, Hisashi Shimoji, Jessica Severin, Takeya Kasukawa, Serkan Sahin, Imad Abugessaisa, Shiro Fukuda, Fumi Hori, Sachi Ishikawa-Kato, Christopher J Mungall, Erik Arner J Kenneth Baillie, Nicolas Bertin, Hidemasa Bono, Michiel de Hoon, Alexander D Diehl, Emmanuel Dimont, Tom C Freeman, Kaori Fujieda, Winston Hide, Rajaram Kaliyaperumal, Toshiaki Katayama, Timo Lassmann, Terrence F Meehan, Koro Nishikata, Hiromasa Ono, Michael Rehli, Albin Sandelin, Erik A Schultes, Peter A C ‘t Hoen, Zuotian Tatum, Mark Thompson, Tetsuro Toyoda, Derek W Wright, Carsten O Daub, Masayoshi Itoh, Piero Carninci, Yoshihide Hayashizaki, Alistair R R Forrest, Hideya Kawaji, and FANTOM consortium. Gateways to the FANTOM5 promoter level mammalian expression atlas. Genome Biology, 16(1):22, 2015. ISSN 1465-6906. doi: 10.1186/s13059-014-0560-6.

17. Raquel García-Pérez, Jose Miguel Ramirez, Aida Ripoll-Cladellas, Ruben Chazarra-Gil, Winona Oliveros, Oleksandra Soldatkina, Mattia Bosio, Paul Joris Rognon, Salvador Capella-Gutierrez, Miquel Calvo, Ferran Reverter, Roderic Guigó, François Aguet, Pedro G. Ferreira, Kristin G. Ardlie, and Marta Melé. The landscape of expression and alternative splicing variation across human traits. Cell Genomics, 3(1):100244, 2023. ISSN 2666-979X. doi: 10.1016/j.xgen.2022.100244.

18. Christina J Herrmann, Ralf Schmidt, Alexander Kanitz, Panu Artimo, Andreas J Gruber, and Mihaela Zavolan. PolyASite 2.0: a consolidated atlas of polyadenylation sites from 3 end sequencing. Nucleic Acids Research, 48(D1):D174–D179, 2019. ISSN 0305-1048. doi: 10.1093/nar/gkz918.

19. Heng Li. Minimap2: pairwise alignment for nucleotide sequences. Bioinformatics, 34(18): 3094–3100, 2018. ISSN 1367-4803. doi: 10.1093/bioinformatics/bty191.

20. Dana Wyman and Ali Mortazavi. TranscriptClean: variant-aware correction of indels, mismatches and splice junctions in long-read transcripts. Bioinformatics, 35(2):340–342, 2019. ISSN 1367-4803. doi: 10.1093/bioinformatics/bty483.

21. Dana Wyman, Gabriela Balderrama-Gutierrez, Fairlie Reese, Shan Jiang, Sorena Rahmanian, Stefania Forner, Dina Matheos, Weihua Zeng, Brian Williams, Diane Trout, Whitney England, Shu-Hui Chu, Robert C. Spitale, Andrea J. Tenner, Barbara J. Wold, and Ali Mortazavi. A technology-agnostic long-read analysis pipeline for transcriptome discovery and quantification. bioRxiv, page 672931, 2020. doi: 10.1101/672931.

22. Muhammed Hasan Çelik and Ali Mortazavi. Analysis of alternative polyadenylation from long-read or short-read RNA-seq with LAPA. bioRxiv, page 2022.11.08.515683, 2022. doi: 10.1101/2022.11.08.515683.

23. Dafni A. Glinos, Garrett Garborcauskas, Paul Hoffman, Nava Ehsan, Lihua Jiang, Alper Gokden, Xiaoguang Dai, François Aguet, Kathleen L. Brown, Kiran Garimella, Tera Bowers, Maura Costello, Kristin Ardlie, Ruiqi Jian, Nathan R. Tucker, Patrick T. Ellinor, Eoghan D. Harrington, Hua Tang, Michael Snyder, Sissel Juul, Pejman Mohammadi, Daniel G. MacArthur, Tuuli Lappalainen, and Beryl B. Cummings. Transcriptome variation in human tissues revealed by long-read sequencing. Nature, pages 1–8, 2022. ISSN 0028-0836. doi: 10.1038/s41586-022-05035-y.

24. Manuel Tardaguila, Lorena de la Fuente, Cristina Marti, Cécile Pereira, Francisco Jose Pardo-Palacios, Hector del Risco, Marc Ferrell, Maravillas Mellado, Marissa Macchietto, Kenneth Verheggen, Mariola Edelmann, Iakes Ezkurdia, Jesus Vazquez, Michael Tress, Ali Mortazavi, Lennart Martens, Susana Rodriguez-Navarro, Victoria Moreno-Manzano, and Ana Conesa. SQANTI: extensive characterization of long-read transcript sequences for quality control in full-length transcriptome identification and quantification. Genome Research, 28(3):396–411, 2018. ISSN 1088-9051. doi: 10.1101/gr.222976.117.

25. Anoushka Joglekar, Andrey Prjibelski, Ahmed Mahfouz, Paul Collier, Susan Lin, Anna Katharina Schlusche, Jordan Marrocco, Stephen R. Williams, Bettina Haase, Ashley Hayes, Jennifer G. Chew, Neil I. Weisenfeld, Man Ying Wong, Alexander N. Stein, Simon A. Hardwick, Toby Hunt, Qi Wang, Christoph Dieterich, Zachary Bent, Olivier Fedrigo, Steven A. Sloan, Davide Risso, Erich D. Jarvis, Paul Flicek, Wenjie Luo, Geoffrey S. Pitt, Adam Frankish, August B. Smit, M. Elizabeth Ross, and Hagen U. Tilgner. A spatially resolved brain region- and cell type-specific isoform atlas of the postnatal mouse brain. Nature Communications, 12(1):463, 2021. doi: 10.1038/s41467-020-20343-5.

26. Alfredo Castello, Bernd Fischer, Katrin Eichelbaum, Rastislav Horos, Benedikt M. Beckmann, Claudia Strein, Norman E. Davey, David T. Humphreys, Thomas Preiss, Lars M. Steinmetz, Jeroen Krijgsveld, and Matthias W. Hentze. Insights into RNA Biology from an Atlas of Mammalian mRNA-Binding Proteins. Cell, 149(6):1393–1406, 2012. ISSN 0092-8674. doi: 10.1016/j.cell.2012.04.031.

27. S B Martins, T Eide, R L Steen, T Jahnsen, S Skalhegg B, and P Collas. HA95 is a protein of the chromatin and nuclear matrix regulating nuclear envelope dynamics. Journal of Cell Science, 113(21):3703–3713, 2000. ISSN 0021-9533. doi: 10.1242/jcs.113.21.3703.

28. Nigel G. Laing, Danielle E. Dye, Carina Wallgren-Pettersson, Gabriele Richard, Nicole Monnier, Suzanne Lillis, Thomas L. Winder, Hanns Lochmüller, Claudio Graziano, Stella Mitrani-Rosenbaum, Darren Twomey, John C. Sparrow, Alan H. Beggs, and Kristen J. Nowak. Mutations and polymorphisms of the skeletal muscle α-actin gene (ACTA1). Human Mutation, 30(9):1267–1277, 2009. ISSN 1059-7794. doi: 10.1002/humu.21059.

29. Sara S. Procknow and Beth A. Kozel. Emerging mechanisms of elastin transcriptional regulation. American Journal of Physiology-Cell Physiology, 323(3):C666–C677, 2022. ISSN 0363-6143. doi: 10.1152/ajpcell.00228.2022.

30. Matthieu Lacroix, Geneviève Rodier, Olivier Kirsh, Thibault Houles, Hélène Delpech, Berfin Seyran, Laurie Gayte, Francois Casas, Laurence Pessemesse, Maud Heuillet, Floriant Bellvert, Jean-Charles Portais, Charlene Berthet, Florence Bernex, Michele Brivet, Audrey Boutron, Laurent Le Cam, and Claude Sardet. E4F1 controls a transcriptional program essential for pyruvate dehydrogenase activity. Proceedings of the National Academy of Sciences, 113(39):10998–11003, 2016. ISSN 0027-8424. doi: 10.1073/pnas.1602754113.

31. R A Kahn, F G Kern, J Clark, E P Gelmann, and C Rulka. Human ADP-ribosylation factors. A functionally conserved family of GTP-binding proteins. Journal of Biological Chemistry, 266(4):2606–2614, 1991. ISSN 0021-9258. doi: 10.1016/s0021-9258(18)52288-2.32.

32. 2013.

33. Michael L. Tress, Federico Abascal, and Alfonso Valencia. Alternative Splicing May Not Be the Key to Proteome Complexity. Trends in Biochemical Sciences, 42(2):98–110, 2017. ISSN 0968-0004. doi: 10.1016/j.tibs.2016.08.008.

34. Julien Lagarde, Barbara Uszczynska-Ratajczak, Silvia Carbonell, Sílvia Pérez-Lluch, Amaya Abad, Carrie Davis, Thomas R Gingeras, Adam Frankish, Jennifer Harrow, Roderic Guigo, and Rory Johnson. High-throughput annotation of full-length long noncoding RNAs with capture long-read sequencing. Nature Genetics, 49(12):1731–1740, 2017. ISSN 1061-4036. doi: 10.1038/ng.3988.

35. Angela Brooks, Francisco Pardo-Palacios, Fairlie Reese, Silvia Carbonell-Sala, Mark Diekhans, Cindy Liang, Dingjie Wang, Brian Williams, Matthew Adams, Amit Behera, Julien Lagarde, Haoran Li, Andrey Prjibelski, Gabriela Balderrama-Gutierrez, Muhammed Hasan Çelik, Maite D. María, Nancy Denslow, Natàlia Garcia-Reyero, Stefan Goetz, Margaret Hunter, Jane Loveland, Carlos Menor, David Moraga, Jonathan Mudge, Hazuki Takahashi, Alison Tang, Ingrid Youngworth, Piero Carninci, Roderic Guigó, Hagen Tilgner, Barbara Wold, Christopher Vollmers, Gloria Sheynkman, Adam Frankish, Kin Fai Au, Ana Conesa, and Ali Mortazavi. Systematic assessment of long-read RNA-seq methods for transcript identification and quantification. 2021. doi: 10.21203/rs.3.rs-777702/v1.

36. Simon A. Hardwick, Wen Hu, Anoushka Joglekar, Li Fan, Paul G. Collier, Careen Foord, Jennifer Balacco, Samantha Lanjewar, Maureen McGuirk Sampson, Frank Koopmans, Andrey D. Prjibelski, Alla Mikheenko, Natan Belchikov, Julien Jarroux, Anne Bergstrom Lucas, Miklós Palkovits, Wenjie Luo, Teresa A. Milner, Lishomwa C. Ndhlovu, August B. Smit, John Q. Trojanowski, Virginia M. Y. Lee, Olivier Fedrigo, Steven A. Sloan, Dóra Tombácz, M. Elizabeth Ross, Erich Jarvis, Zsolt Boldogkö i, Li Gan, and Hagen U. Tilgner. Singlenuclei isoform RNA sequencing unlocks barcoded exon connectivity in frozen brain tissue. Nature Biotechnology, 40(7):1082–1092, 2022. ISSN 1087-0156. doi: 10.1038/s41587-022-01231-3.

37. Elisabeth Rebboah, Fairlie Reese, Katherine Williams, Gabriela Balderrama-Gutierrez, Cassandra McGill, Diane Trout, Isaryhia Rodriguez, Heidi Liang, Barbara J. Wold, and Ali Mortazavi. Mapping and modeling the genomic basis of differential RNA isoform expression at single-cell resolution with LR-Split-seq. Genome Biology, 22(1):286, 2021. doi: 10.1186/s13059-021-02505-w.

38. Alison D. Tang, Cameron M. Soulette, Marijke J. van Baren, Kevyn Hart, Eva Hrabeta-Robinson, Catherine J. Wu, and Angela N. Brooks. Full-length transcript characterization of SF3B1 mutation in chronic lymphocytic leukemia reveals downregulation of retained introns. Nature Communications, 11(1):1438, 2020. doi: 10.1038/s41467-020-15171-6.

39. Feng Yue, Yong Cheng, Alessandra Breschi, Jeff Vierstra, Weisheng Wu, Tyrone Ryba, Richard Sandstrom, Zhihai Ma, Carrie Davis, Benjamin D Pope, Yin Shen, Dmitri D Pervouchine, Sarah Djebali, Robert E Thurman, Rajinder Kaul, Eric Rynes, Anthony Kirilusha, Georgi K Marinov, Brian A Williams, Diane Trout, Henry Amrhein, Katherine Fisher-Aylor, Igor Antoshechkin, Gilberto DeSalvo, Lei-Hoon See, Meagan Fastuca, Jorg Drenkow, Chris Zaleski, Alex Dobin, Pablo Prieto, Julien Lagarde, Giovanni Bussotti, Andrea Tanzer, Olgert Denas, Kanwei Li, M A Bender, Miaohua Zhang, Rachel Byron, Mark T Groudine, David McCleary, Long Pham, Zhen Ye, Samantha Kuan, Lee Edsall, Yi-Chieh Wu, Matthew D Rasmussen, Mukul S Bansal, Manolis Kellis, Cheryl A Keller, Christapher S Morrissey, Tejaswini Mishra, Deepti Jain, Nergiz Dogan, Robert S Harris, Philip Cayting, Trupti Kawli, Alan P Boyle, Ghia Euskirchen, Anshul Kundaje, Shin Lin, Yiing Lin, Camden Jansen, Venkat S Malladi, Melissa S Cline, Drew T Erickson, Vanessa M Kirkup, Katrina Learned, Cricket A Sloan, Kate R Rosenbloom, Beatriz Lacerda de Sousa, Kathryn Beal, Miguel Pignatelli, Paul Flicek, Jin Lian, Tamer Kahveci, Dongwon Lee W James Kent, Miguel Ramalho Santos, Javier Herrero, Cedric Notredame, Audra Johnson, Shinny Vong, Kristen Lee, Daniel Bates, Fidencio Neri, Morgan Diegel, Theresa Canfield, Peter J Sabo, Matthew S Wilken, Thomas A Reh, Erika Giste, Anthony Shafer, Tanya Kutyavin, Eric Haugen, Douglas Dunn, Alex P Reynolds, Shane Neph, Richard Humbert R Scott Hansen, Marella De Bruijn, Licia Selleri, Alexander Rudensky, Steven Josefowicz, Robert Samstein, Evan E Eichler, Stuart H Orkin, Dana Levasseur, Thalia Papayannopoulou, Kai-Hsin Chang, Arthur Skoultchi, Srikanta Gosh, Christine Disteche, Piper Treuting, Yanli Wang, Mitchell J Weiss, Gerd A Blobel, Xiaoyi Cao, Sheng Zhong, Ting Wang, Peter J Good, Rebecca F Lowdon, Leslie B Adams, Xiao-Qiao Zhou, Michael J Pazin, Elise A Feingold, Barbara Wold, James Taylor, Ali Mortazavi, Sherman M Weissman, John A Stamatoyannopoulos, Michael P Snyder, Roderic Guigo, Thomas R Gingeras, David M Gilbert, Ross C Hardison, Michael A Beer, Bing Ren, and The Mouse ENCODE Consortium. A comparative encyclopedia of DNA elements in the mouse genome. Nature, 515(7527):355–364, 2014. ISSN 0028-0836. doi: 10.1038/nature13992.

40. Hazuki Takahashi, Sachi Kato, Mitsuyoshi Murata, and Piero Carninci. Gene Regulatory Networks, Methods and Protocols. Methods in Molecular Biology, 786:181–200, 2011. ISSN 1064-3745. doi: 10.1007/978-1-61779-292-2\_11.

41. Philippe Batut and Thomas R. Gingeras. RAMPAGE: Promoter Activity Profiling by Paired-End Sequencing of 5-Complete cDNAs. Current Protocols in Molecular Biology, 104(1):p 25B.11.1–25B.11.16, 2013. ISSN 1934-3639. doi: 10.1002/0471142727.mb25b11s104.

42. Lingyun Song and Gregory E. Crawford. DNase-seq: A High-Resolution Technique for Mapping Active Gene Regulatory Elements across the Genome from Mammalian Cells. Cold Spring Harbor Protocols, 2010(2):ppdb.prot5384, 2010. ISSN 1940-3402. doi: 10.1101/pdb.prot5384.

43. Juan L. Trincado, Juan C. Entizne, Gerald Hysenaj, Babita Singh, Miha Skalic, David J. Elliott, and Eduardo Eyras. SUPPA2: fast, accurate, and uncertainty-aware differential splicing analysis across multiple conditions. Genome Biology, 19(1):40, 2018. ISSN 1474-7596. doi: 10.1186/s13059-018-1417-1.

44. Jose Manuel Rodriguez, Paolo Maietta, Iakes Ezkurdia, Alessandro Pietrelli, Jan-Jaap Wesselink, Gonzalo Lopez, Alfonso Valencia, and Michael L. Tress. APPRIS: annotation of principal and alternative splice isoforms. Nucleic Acids Research, 41(D1):D110–D117, 2013. ISSN 0305-1048. doi: 10.1093/nar/gks1058.

45. Dominik M. Endres and Johannes E. Schindelin. A New Metric for Probability Distributions. IEEE Transactions on Information Theory, 49(7):1858, 2003. ISSN 0018-9448. doi: 10.1109/tit.2003.813506.

46. Pauli Virtanen, Ralf Gommers, Travis E Oliphant, Matt Haberland, Tyler Reddy, David Cournapeau, Evgeni Burovski, Pearu Peterson, Warren Weckesser, Jonathan Bright, Stéfan J van der Walt, Matthew Brett, Joshua Wilson, K Jarrod Millman, Nikolay Mayorov, Andrew R J Nelson, Eric Jones, Robert Kern, Eric Larson, C J Carey, ?Ilhan Polat, Yu Feng, Eric W Moore, Jake VanderPlas, Denis Laxalde, Josef Perktold, Robert Cimrman, Ian Henriksen, E A Quintero, Charles R Harris, Anne M Archibald, Antônio H Ribeiro, Fabian Pedregosa, Paul van Mulbregt, SciPy 1 0 Contributors, Aditya Vijaykumar, Alessandro Pietro Bardelli, Alex Rothberg, Andreas Hilboll, Andreas Kloeckner, Anthony Scopatz, Antony Lee, Ariel Rokem, C Nathan Woods, Chad Fulton, Charles Masson, Christian Häggström, Clark Fitzgerald, David A Nicholson, David R Hagen, Dmitrii V Pasechnik, Emanuele Olivetti, Eric Martin, Eric Wieser, Fabrice Silva, Felix Lenders, Florian Wilhelm, G Young, Gavin A Price, Gert-Ludwig Ingold, Gregory E Allen, Gregory R Lee, Hervé Audren, Irvin Probst, Jörg P Dietrich, Jacob Silterra, James T Webber, Janko Slavić, Joel Nothman, Johannes Buchner, Johannes Kulick, Johannes L Schönberger, José Vinícius de Miranda Cardoso, Joscha Reimer, Joseph Harrington, Juan Luis Cano Rodríguez, Juan Nunez-Iglesias, Justin Kuczynski, Kevin Tritz, Martin Thoma, Matthew Newville, Matthias Kümmerer, Maximilian Bolingbroke, Michael Tartre, Mikhail Pak, Nathaniel J Smith, Nikolai Nowaczyk, Nikolay Shebanov, Oleksandr Pavlyk, Per A Brodtkorb, Perry Lee, Robert T McGibbon, Roman Feldbauer, Sam Lewis, Sam Tygier, Scott Sievert, Sebastiano Vigna, Stefan Peterson, Surhud More, Tadeusz Pudlik, Takuya Oshima, Thomas J Pingel, Thomas P Robitaille, Thomas Spura, Thouis R Jones, Tim Cera, Tim Leslie, Tiziano Zito, Tom Krauss, Utkarsh Upadhyay, Yaroslav O Halchenko, and Yoshiki Vázquez-Baeza. SciPy 1.0: fundamental algorithms for scientific computing in Python. Nature Methods, 17(3):261–272, 2020. ISSN 1548-7091. doi: 10.1038/s41592-019-0686-2.

47. Fairlie Reese and Ali Mortazavi. Swan: a library for the analysis and visualization of longread transcriptomes. Bioinformatics, 37(9):btaa836–, 2020. ISSN 1367-4803. doi: 10.1093/bioinformatics/btaa836.

48. Richard I. Kuo, Yuanyuan Cheng, Runxuan Zhang, John W. S. Brown, Jacqueline Smith, Alan L. Archibald, and David W. Burt. Illuminating the dark side of the human transcriptome with long read transcript sequencing. BMC Genomics, 21(1):751, 2020. doi: 10.1186/s12864-020-07123-7.

49. Stephen F. Altschul, Warren Gish, Webb Miller, Eugene W. Myers, and David J. Lipman. Basic local alignment search tool. Journal of Molecular Biology, 215(3):403–410, 1990. ISSN 0022-2836. doi: 10.1016/s0022-2836(05)80360-2.

50. Aaron R. Quinlan and Ira M. Hall. BEDTools: a flexible suite of utilities for comparing genomic features. Bioinformatics, 26(6):841–842, 2010. ISSN 1367-4803. doi: 10.1093/bioinformatics/btq033.

